# CorB and MgpB (YegH) alleviate excess-magnesium stress and contribute to egg white tolerance and virulence in *Salmonella*

**DOI:** 10.64898/2026.09.30.755624

**Authors:** Yumi Iwadate, James M. Slauch

## Abstract

Maintaining proper cation homeostasis is essential for the survival of all living organisms. During infection, *Salmonella* must adapt to host-imposed magnesium (Mg²⁺) limitation. To compensate, *Salmonella* induces high-affinity Mg²⁺ transporters and synthesis of polyamines, organic cations that partially substitute for Mg²⁺. Once Mg²⁺ levels are restored and cells enter stationary phase, excess cations must be exported; failure to do so results in loss of viability. The CorC-domain protein PaeA exports excess polyamines, whereas MgpA and CorC reduce intracellular Mg²⁺ levels. Here, we identify two new CorC-domain proteins, MgpB (formerly YegH) and CorB, as mediators of Mg²⁺ homeostasis, showing that they also reduce intracellular Mg²⁺ accumulation and protect cells from endogenous excess-cation stress. Genetic analyses revealed the functional redundancy among these proteins in conferring resistance to exogenous high Mg²⁺, endogenous stress induced after Mg²⁺ starvation, and egg whites. Deletion of *mgpB* and/or *corB* conferred little to no phenotype. However, these mutations showed synergistic phenotypes in strains lacking MgpA. This synergistic relationship was also evident in mouse competition assays. Lack of MgpA, CorC, MgpB, and CorB leads to an increase in intracellular Mg²⁺ levels. Moreover, expression of *mgpB* and *corB* are increased in high Mg²⁺, all consistent with their role as putative Mg²⁺ exporters. Together, these findings identify MgpB and CorB as new mediators of Mg²⁺ homeostasis and reveal a multilayered system that enables *Salmonella* to finely balance intracellular cation levels during environmental and host-imposed stress.

**IMPORTANCE:** Mg²⁺ neutralizes negative charges within cells and influences macromolecular structure, stability, function, and intermolecular interactions. Mg²⁺ cannot be synthesized or degraded, requiring tight regulation through coordinated influx and efflux. However, the mechanisms that downregulate Mg²⁺ levels remain poorly understood. In this work, we identify two CorC-domain proteins, MgpB and CorB, as critical components of Mg²⁺ homeostasis that protect cells from both endogenous and exogenous excess-Mg^2+^ stress. Importantly, these proteins also contribute to virulence and egg white tolerance, highlighting cation homeostasis as a key target of host antimicrobial defenses. These findings and previous work establish the CorC-domain family of proteins as a layered, partially redundant system for maintaining cation balance.

## INTRODUCTION

Magnesium (Mg²⁺) is an essential element for all living organisms. As a cofactor, Mg²⁺ is required for numerous enzymatic reactions, including many involving ATP hydrolysis (1, 2). As a hydrated divalent cation, Mg²⁺ neutralizes negative charges within the cell, including phosphate groups in DNA, RNA, ribosomes, and lipopolysaccharides (3–7). Thus, Mg²⁺ homeostasis is central to cell viability. While Mg²⁺ import systems have been extensively characterized (8, 9), Mg²⁺ export remains poorly understood. CorA, the primary Mg²⁺ importer under Mg²⁺-replete conditions, has been reported to mediate Mg²⁺ efflux, although the physiological significance of this activity remains unclear (10). More recently, CorB proteins were shown to function as Mg²⁺ efflux transporters in thermophilic archaea and bacteria (11, 12), and mutations in *corB* homologs in Gram-positive bacteria confer sensitivity to high Mg²⁺ and increased intracellular Mg²⁺ accumulation (13, 14). In contrast, the mechanisms of Mg²⁺ export in Gram-negative bacteria remain poorly understood, despite the presence of CorB homologs in model organisms such as *Escherichia coli,* likely because their deletion produces no readily detectable phenotype. Our previous work identified MgpA and CorC as major contributors to reducing intracellular Mg²⁺ levels in *S.* Typhimurium (15).

*Salmonella* is a foodborne pathogen that causes not only gastroenteritis but also severe systemic disease and sepsis (16). A key feature of *Salmonella* pathogenesis is its ability to survive and replicate within macrophage phagosomes, an environment that normally restricts bacterial growth (17–19). Macrophages limit intracellular bacterial proliferation through multiple mechanisms, including phagosomal acidification (20, 21) and restriction of Mg²⁺ via the phagosomal membrane transporter SLC11A1 (NRAMP1)(19, 22). In response, *Salmonella* activates the PhoPQ two-component system, a master regulator of the low-Mg²⁺ stress response (23, 24) that regulates numerous genes (25, 26), including genes encoding high-affinity Mg²⁺ transporters, lipopolysaccharide modification enzymes, and additional virulence factors (27–29).

We previously showed that Mg²⁺ starvation induces, not only the high-affinity Mg²⁺ transporters MgtA and MgtB, but also the synthesis of polyamines (30, 31). We hypothesize that these organic cationic polyamines partially substitute for Mg²⁺ under Mg²⁺ limiting conditions (32). Either Mg²⁺ transport or polyamine synthesis is required to maintain viability under low-Mg²⁺ conditions (30) (Fig.1). Counterintuitively, upon entry into stationary phase, these compensatory pathways lead to a lethal overload of cations. At stationary phase, excess polyamines must be exported by the inner-membrane transporter PaeA (30), while excess Mg²⁺ is reduced by MgpA and CorC (15); loss of either system results in cell death during stationary phase (Fig.1). Importantly, these lethal phenotypes are suppressed by blocking Mg²⁺ transport or polyamine synthesis, demonstrating that the accumulation of Mg²⁺ and polyamines, termed excess-cation stress, is toxic (15, 30). Consistent with these functions, PaeA, but not MgpA and CorC, is required for tolerance to high polyamine stress (33), and MgpA and CorC, but not PaeA, are required for tolerance to high Mg²⁺ stress (15). Thus, PaeA versus MgpA and CorC represent two distinct but coordinated cation-control systems. This phenomenon is recapitulated during infection, supporting a model in which *Salmonella* must coordinately regulate Mg²⁺ and polyamine levels to maintain cation homeostasis within the macrophage phagosome (15, 30).

**Fig. 1.**
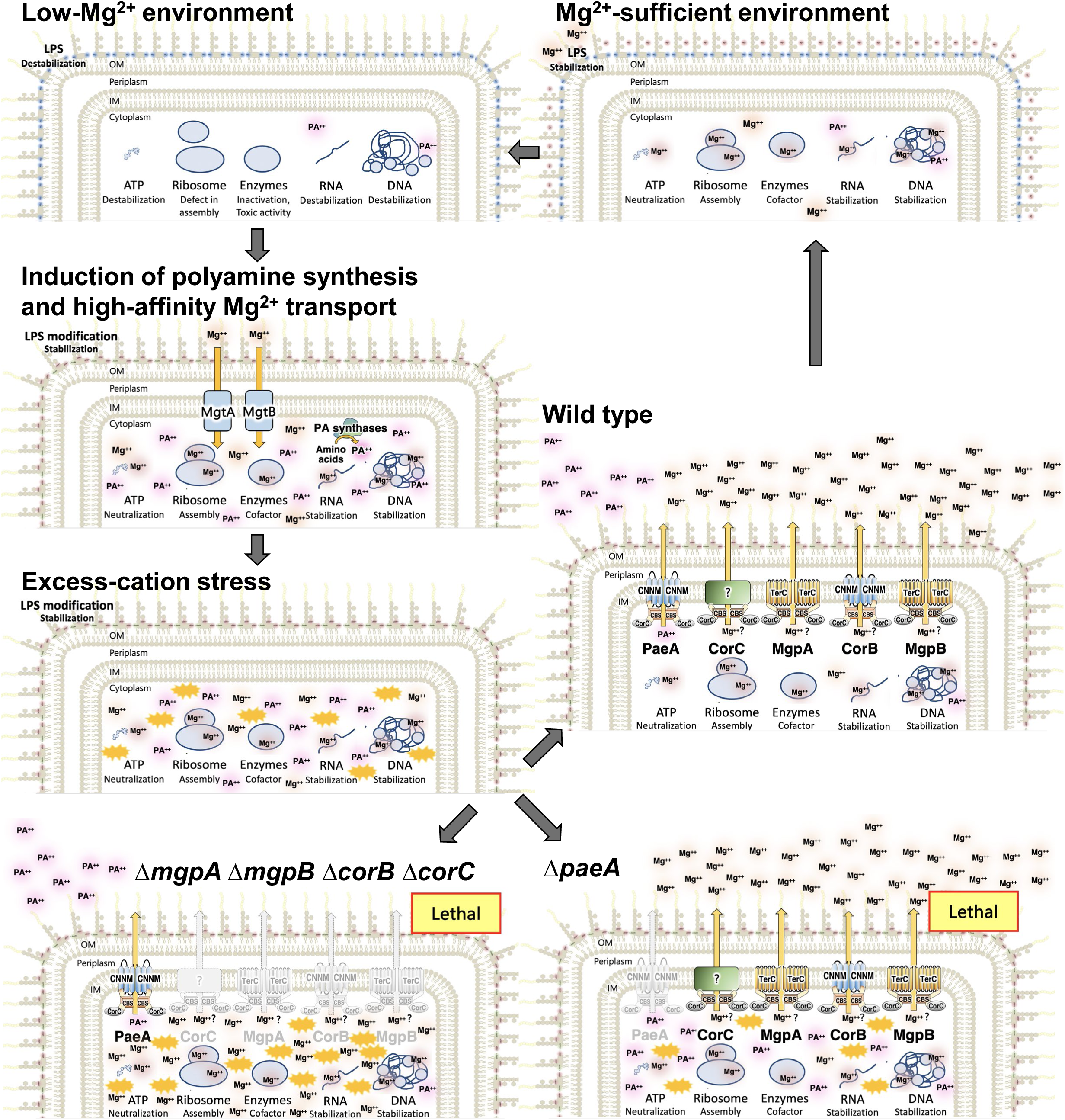
Excess-cation stress in *Salmonella.* Under Mg²⁺ starvation, *Salmonella* initially experiences low-cation stress, which induces the high-affinity Mg²⁺ transporters MgtA and MgtB to scavenge trace Mg²⁺ from the environment, as well as polyamine synthesis to compensate for Mg²⁺ deficiency. Upon entry to stationary phase, endogenous excess-cation stress arises. The CorC-domain-containing proteins PaeA and MgpA/MgpB (YegH)/CorB/CorC alleviate this stress by reducing intracellular polyamine and Mg²⁺ levels, respectively.

The CorC protein domain (Pfam03471) is widespread throughout Bacteria and Archaea but remains functionally uncharacterized (34). Notably, all proteins identified as important for survival during both endogenous and exogenous excess-cation stress, PaeA, MgpA, and CorC, contain a CorC domain. The *Salmonella* genome encodes six proteins containing CorC domains: PaeA (YtfL), CorB, CorC, MgpA (YoaE), MgpB (YegH), and CvrA. These proteins each contain cytoplasmic CBS pair and CorC domains, but differ in their N-terminal transmembrane architectures, suggesting functional specialization (Fig. 2). PaeA consists of an N-terminal CNNM transmembrane domain (pfam01595), a cytoplasmic CBS-pair domain (pfam00571), and a C-terminal CorC domain. CorB, which is a homolog of known Mg²⁺ efflux transporters (11, 12), resembles PaeA in domain architecture, containing a CNNM transmembrane domain, a CBS pair that binds Mg²⁺-ATP at the dimer interface (12), and a CorC domain. The eponymous CorC lacks a transmembrane domain and consists only of a CBS pair and CorC domain. MgpA and MgpB share a distinct architecture, with an N-terminal TerC transmembrane domain (pfam03741), followed by a CBS pair and CorC domain. CvrA contains an N-terminal Na⁺/H⁺ exchanger domain (pfam00999), a cytoplasmic RCK domain (pfam02080), and a C-terminal CorC domain. CvrA in *Vibrio cholerae* functions as a K⁺/H⁺ exchanger (35). Notably, all characterized CorC-domain– containing proteins (CorC proteins) are implicated in cation transport. Therefore, we speculate that the CorC domain, together with the associated CBS-pair domain, functions as a sensor of cytoplasmic cations such as Mg²⁺ (12, 36).

**Fig. 2.**
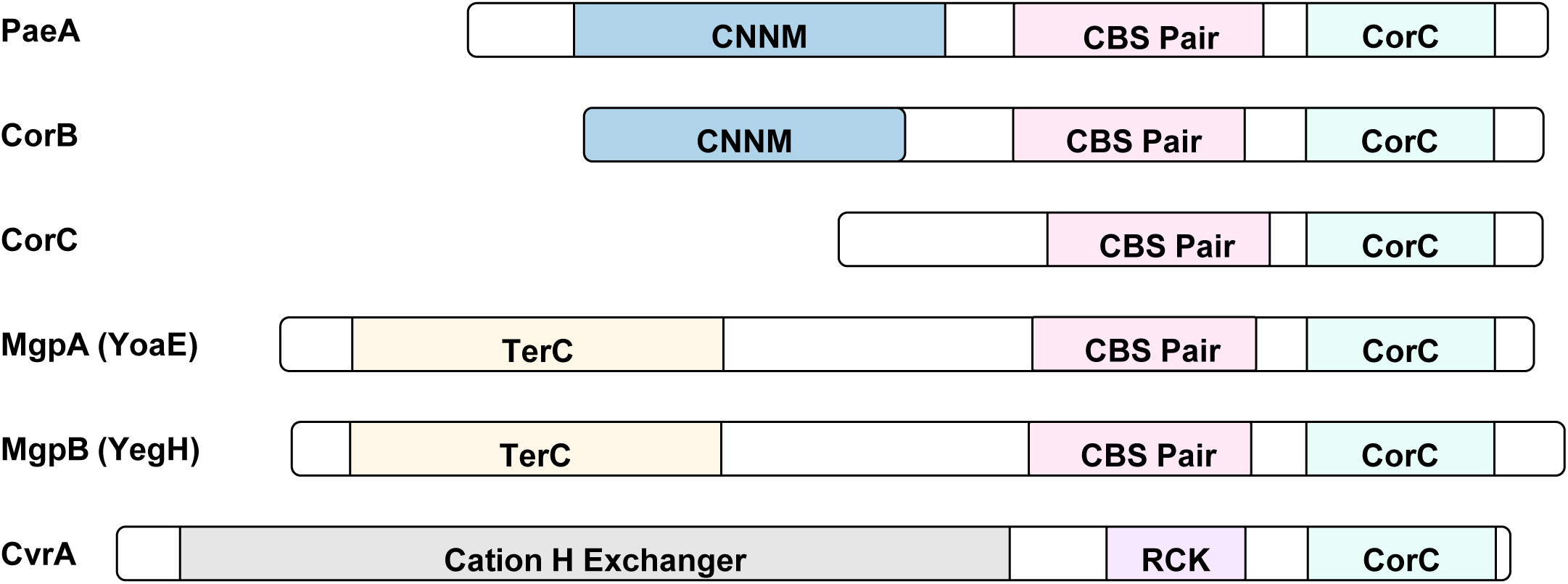
Domain architecture of CorC domain-containing proteins in *Salmonella*. Conserved domains were predicted using InterPro (https://www.ebi.ac.uk/interpro/). Each domain is represented by a different color, while regions with no domain assignment are shown in white. Adapted from (15).

In this study, we extend this framework by identifying two additional CorC-domain-containing proteins, CorB and YegH (renamed MgpB; Mg²⁺ protection protein B), as key contributors to Mg²⁺ efflux and Mg²⁺ homeostasis in *Salmonella*. MgpB shares the same domain architecture with MgpA. In the absence of MgpA, deletion of *mgpB and corB* causes a drastic loss of viability in high Mg²⁺, comparable to that of the Δ*mgpA* Δ*corC* mutant (15), and partially suppresses the sensitivity of a *paeA* strain to high polyamines. The *mgpA mgpB corB* deletion strain accumulates elevated levels of intracellular Mg²⁺. Moreover, *mgpB* and *corB* expression is induced when grown with high Mg²⁺ levels, further suggesting that MgpB and CorB function specifically in Mg²⁺ homeostasis. We also show that MgpB and CorB are required for full virulence and egg white tolerance. Together, our findings define the role of MgpB and CorB in Mg²⁺ homeostasis (Fig.1), acting redundantly with MgpA and CorC, but distinct from the polyamine-specific PaeA exporter. This reveals a multilayered strategy by which *Salmonella* maintains cation homeostasis in response to environmental changes, including during host infection.

## RESULTS

### MgpB and CorB are required for tolerance to exogenous excess-Mg²⁺ stress

We previously demonstrated that the CorC proteins MgpA and CorC function redundantly to confer tolerance to exogenous excess-Mg²⁺ stress, as the Δ*mgpA* Δ*corC* double mutant, but neither single mutant, is sensitive to high Mg²⁺ under alkaline conditions (pH 8.5) (15). The role of high pH in this sensitivity is not clear. To determine whether additional CorC proteins contribute to tolerance, we compared the sensitivity to high-Mg²⁺ stress of the wild-type, the Δ*mgpA* Δ*corC* strain, and a strain lacking all CorC-family proteins (ΔCorC6). Cells were grown to mid-exponential phase in N-minimal medium (pH 7.4) containing 1 mM MgCl₂, washed, diluted into N-minimal medium (pH 8.5) supplemented with 1 or 5 mM MgCl₂, and incubated at 37°C. Optical density at 600 nm (OD₆₀₀) was measured after 24 h. In 5 mM MgCl₂, the Δ*mgpA* Δ*corC* mutant showed a mild but reproducible growth defect. The ΔCorC6 strain exhibited substantially greater sensitivity to high Mg²⁺ than the Δ*mgpA* Δ*corC* mutant (Fig. 3A), indicating that additional CorC-family proteins contribute to tolerance of exogenous high-Mg²⁺ stress.

**Fig. 3.**
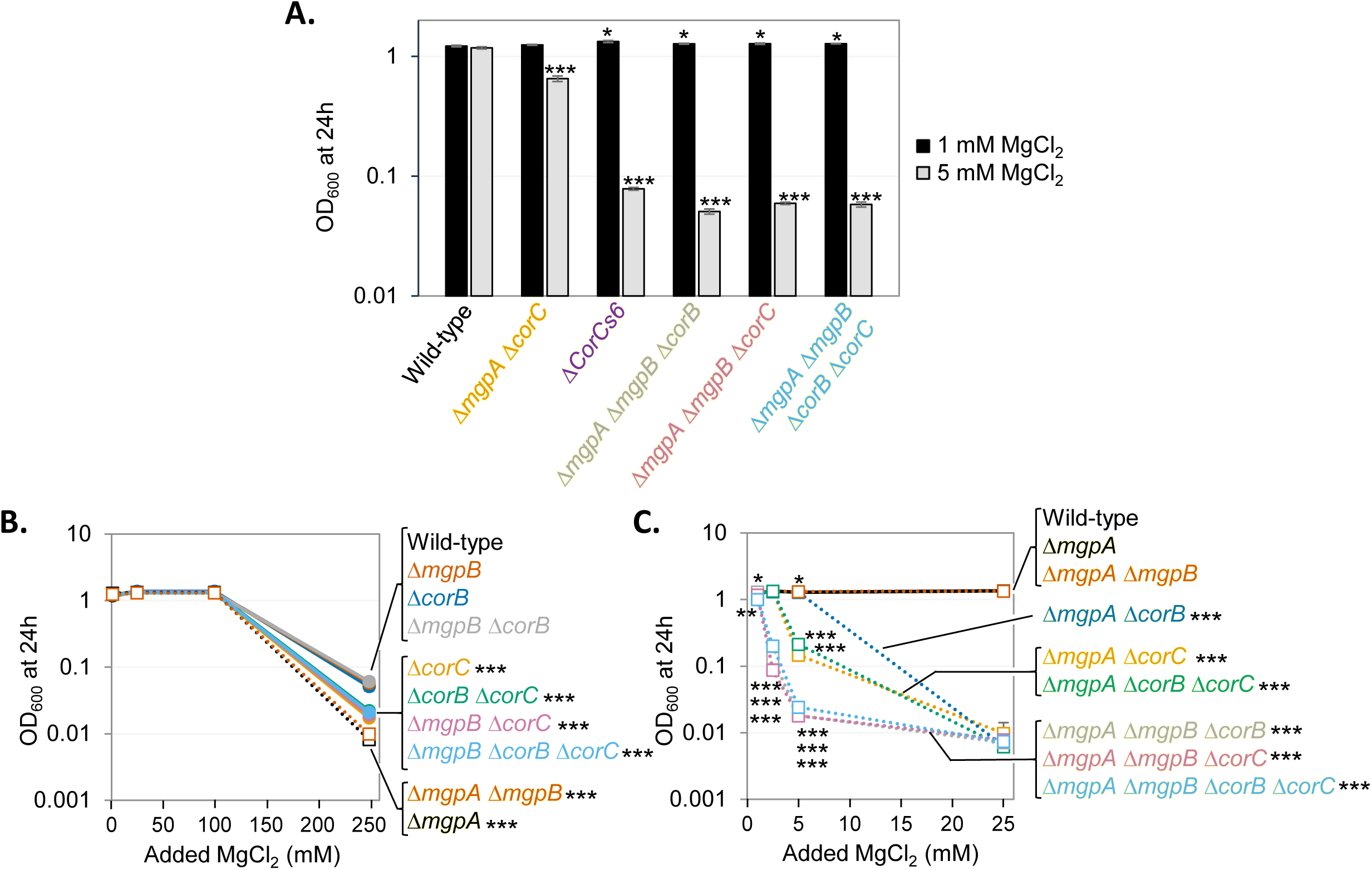
Loss of MgpB or CorB sensitizes cells to exogenous high Mg²⁺ at high pH in the absence of MgpA. The indicated strains were pre-grown to mid-exponential phase in N-minimal medium pH 7.4 with 1 mM MgCl_2_, washed, and diluted into N-minimal medium pH 8.5 with the indicated amount of MgCl_2_, followed by incubation at 37 ℃. OD₆₀₀ was measured after 24 hours of incubation. Values are mean ± SD, n = 6. Unpaired t test (p < 0.05*, 0.005**, 0.0005***) versus corresponding WT at the same condition. *p-*values at 1, 25, and 100 mM MgCl_2_ in panel B are not shown. Complete *p*-values for panels B and C are provided in Tables S4 and S5, respectively. Strains used: 14028, JS2730, JS2736, JS2737, JS2738, JS2739, JS2728, JS2740, JS2741, JS2707, JS2742, JS2743, JS2744, JS2729, JS2745, JS2746, and JS2747.

Using strains carrying different combinations of deletions in CorC protein-encoding genes, we identified the Δ*mgpA* Δ*mgpB* Δ*corB*, Δ*mgpA* Δ*mgpB* Δ*corC*, and Δ*mgpA* Δ*mgpB* Δ*corB* Δ*corC* mutants as highly sensitive to high-Mg²⁺ stress, comparable to that of the ΔCorC6 strain (Fig. 3A). These findings reveal previously unrecognized roles for MgpB and CorB in tolerance to exogenous high-Mg²⁺ stress under alkaline conditions.

To define the functional relationships among *mgpA*, *mgpB*, *corB*, and *corC*, we examined high-Mg²⁺ sensitivity across all combinations of deletions in these genes. In an *mgpA*⁺ background, deletion of *corC*, *corB*, or *mgpB* alone did not affect sensitivity to high Mg²⁺ (Fig. 3B). Deletion of *mgpA* or both *mgpA* and *mgpB* caused only a modest increase in sensitivity, detectable only at the highest MgCl₂ concentration tested (250 mM) (Fig. 3B). In contrast, in the Δ*mgpA* background, deletion of *corC* significantly increased sensitivity to high-Mg²⁺ stress, consistent with our previous report (15) (Fig. 3C). Deletion of *corB* in the Δ*mgpA* background also increased sensitivity, though less severely than deletion of *corC* (Fig. 3C). Notably, deletion of *corB* in the Δ*mgpA* Δ*corC* strain did not further enhance sensitivity, suggesting that *corB* and *corC* exhibit an epistatic relationship. Deletion of *mgpB* alone in the Δ*mgpA* background did not alter sensitivity (Fig. 3B and C). However, deletion of *mgpB* in the Δ*mgpA* Δ*corB*, Δ*mgpA* Δ*corC*, or Δ*mgpA* Δ*corB* Δ*corC* backgrounds markedly increased sensitivity, with all strains exhibiting similar levels of sensitivity (Fig. 3C). These results indicate that *mgpB* displays synthetic interactions with *mgpA* and either *corB* or *corC*. Together, these data show that deletion of *corC* or *corB* sensitizes Δ*mgpA* cells to high-Mg²⁺ stress, and that this sensitivity is further exacerbated by loss of MgpB. These results suggest that MgpA and MgpB, which share a similar domain architecture, act in a partially redundant manner to protect cells from exogenous high-Mg²⁺ stress.

To confirm that reduced OD₆₀₀ reflected loss of viability, we measured CFU at 6.5 h after transfer to high-Mg²⁺ alkaline medium. The effects on CFU were even more dramatic, but the overall trends and conclusions were consistent with OD₆₀₀ data alone (Fig. S1A and B). Furthermore, introduction of plasmid-borne *mgpB* or *corB* into the Δ*mgpA* Δ*mgpB* Δ*corB* strain restored tolerance to high-Mg²⁺ stress (Fig. S2), confirming that loss of *mgpB* and *corB* underlies the observed sensitivity. Collectively, these results establish MgpB and CorB as previously unrecognized contributors to tolerance of exogenous high-Mg²⁺ stress under alkaline conditions.

### The Δ*mgpA* Δ*mgpB* Δ*corB* mutant accumulates Mg^2+^ in the cell

We previously demonstrated that the Δ*mgpA* Δ*corC* strain accumulates higher cellular Mg²⁺ levels than the wild type when grown at neutral pH in medium supplemented with 1 or 10 mM MgCl₂, conditions under which the mutant remains viable (15). MgpB shares a similar domain architecture with MgpA, containing an N-terminal TerC transmembrane domain, which has been implicated in tellurite resistance (37, 38) and manganese transport (39). CorB resembles the polyamine exporter PaeA (33) and is a homolog of known Mg²⁺ efflux transporters (11, 12). Therefore, we hypothesized that MgpB and CorB function as transporters that confer resistance to exogenous high-Mg²⁺ stress by reducing Mg²⁺ levels in the cell.

To test this hypothesis, we employed an established bioassay to assess relative intracellular Mg²⁺ levels (15). In this assay, lysates prepared from strains of interest were tested for their ability to stimulate the growth of a Δ*mgtA* Δ*mgtB* strain in N-minimal medium lacking added Mg²⁺. This method enables sensitive estimation of Mg²⁺ levels in cell lysates using a calibration curve generated by measuring the growth yield of the Δ*mgtA* Δ*mgtB* strain in response to exogenously added MgCl₂. Increasing MgCl₂ concentrations increased growth yield in a concentration-dependent manner, confirming the sensitivity of the assay (Fig. S3A and B). The test strains were grown in N-minimal medium (pH 7.4) supplemented with 10 mM MgCl₂. Importantly, none of the strains exhibited a loss of viability under these neutral-pH conditions (Fig. S3C). After 24 h of incubation, cells were harvested, washed, and lysates were prepared in Chelex-treated PBS. The growth-stimulating activity of the lysates was quantified by measuring the final OD₆₀₀ of an Δ*mgtA* Δ*mgtB* strain after 15 h of incubation in N-minimal medium without added MgCl_2_ (Fig. S3D). Cellular Mg²⁺ levels were then estimated using calibration curves generated as described in the Materials and Methods. As a control, lysates had no effect when 1 mM MgCl₂ was supplemented in the assay (Fig. S3E), indicating that the growth-stimulating activity of the lysates was attributable to their Mg²⁺ content rather than to other components.

In the *mgpA*⁺ background, when grown with 10 mM MgCl₂, strains deleted for *mgpB*, *corC*, or *corB,* either individually or in combination, had intracellular Mg²⁺ levels equivalent to wild type (Fig. 4A). In contrast, in the Δ*mgpA* background, deletion of *corC* markedly increased intracellular Mg²⁺ levels, consistent with our previous report (15), whereas deletion of *mgpB* or *corB* alone had no detectable effect (Fig. 4B). However, simultaneous deletion of *mgpB* and *corB* in the Δ*mgpA* background increased intracellular Mg²⁺ levels (Fig. 4B). Additional deletion of *corC* in the Δ*mgpA* Δ*mgpB* Δ*corB* strain resulted in an additional, albeit modest, increase in intracellular Mg²⁺ accumulation (Fig. 4A and B). These results indicate that *mgpB* and *corB*, like *mgpA* and *corC*, contribute to reducing intracellular Mg²⁺ levels under high-Mg²⁺ conditions. Collectively, these results support our hypothesis that MgpB and CorB contribute to resistance to exogenous high-Mg²⁺ stress by reducing intracellular Mg²⁺ levels.

**Fig. 4.**
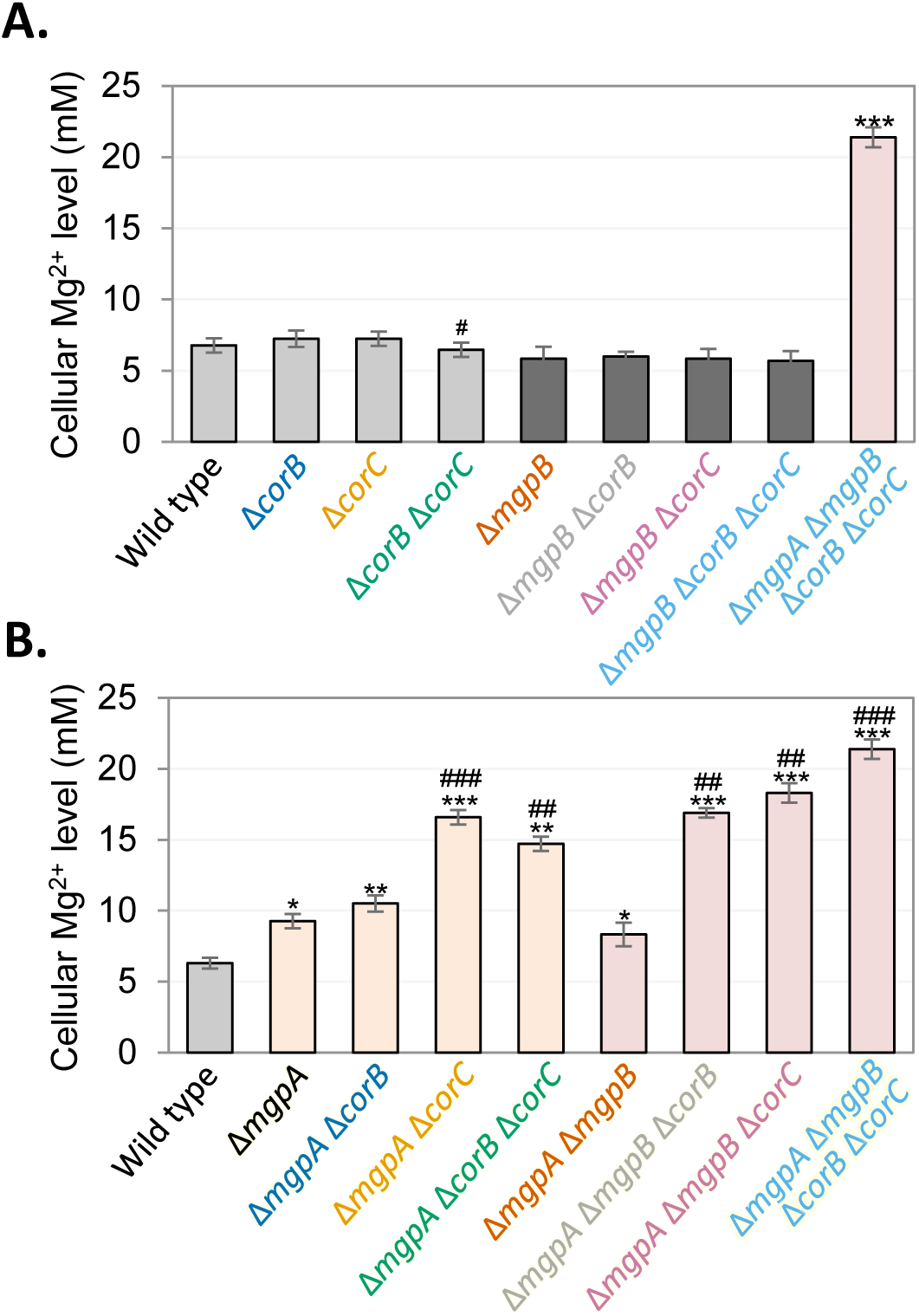
The Δ*mgpA* Δ*mgpB* Δ*corB* strain accumulates Mg². (A and B) Indicated strains were grown in the presence of 10 mM MgCl₂ for 24 h. Cellular Mg²⁺ levels were determined using a bioassay, as described in the Materials and Methods. Briefly, cell lysates were prepared from the indicated strains, and their Mg²⁺ content was estimated based on their ability to stimulate the growth of the Δ*mgtA* Δ*mgtB* strain in medium lacking added Mg²⁺. Levels were calculated from a calibration curve as described. Values are mean ± SD, n = 3. Unpaired t-test (P < 0.05*, 0.005**, 0.0005***) vs. wild type; and (P < 0.05^#^, 0.005^##^, 0.0005^###^) vs. the corresponding *corB*^+^ *corC*^+^ strains. Strains used: JS2562, 14028, JS2728, JS2740, JS2741, JS2730, JS2729, JS2746, JS2745, JS2748, JS2749, JS2750, JS2751, JS2752, JS2753, JS2754, and JS2755.

### MgpB and CorB are required for survival under endogenous excess-cation stress at stationary phase after Mg²⁺ starvation

Our previous work demonstrated that the CorC proteins MgpA and CorC play critical roles in mitigating endogenous excess-cation stress that arises upon entry into stationary phase after Mg²⁺ starvation (15). MgpA and CorC, which function to reduce cellular Mg²⁺ levels, together with the PaeA that reduces cellular divalent polyamines, are required to alleviate this stress; in the absence of these systems, cells lose viability at the transition to stationary phase (15, 30). Although the molecular mechanisms underlying the generation of endogenous excess-cation stress remain unclear, the phenotype appears to result from the combined effects of increased polyamine synthesis and induction of high-affinity Mg²⁺ transport, as deletion of either pathway suppresses the loss of viability in Δ*mgpA* Δ*corC* or Δ*paeA* strains (15, 30). These observations lead to the conclusion that overcompensation in cation accumulation causes toxicity during stationary phase.

Because MgpB and CorB also function to reduce Mg²⁺ levels, similar to MgpA and CorC, we examined whether they are required to alleviate endogenous excess-cation stress during stationary phase after Mg²⁺ starvation. When grown in N-minimal medium (pH 7.4) supplemented with 1 mM MgCl₂, all strains tested grew similarly (Fig. S4A and B) and remained viable throughout 48 h of incubation (Fig. 5A and B). When grown in N-minimal medium (pH 7.4) without added Mg²⁺, growth was comparable among all strains during exponential phase (Fig. S4C and D). However, upon entry into stationary phase, the *corC* deletion strain lost viability, as we previously reported (15), whereas deletion of *mgpB*, *corB*, or both had no effect (Fig. 5C). In the Δ*mgpA* background, deletion of *corC* caused a more severe loss of viability upon transition to stationary phase, consistent with prior observations (15) (Fig. 5D). Deletion of either *mgpB* or *corB* alone did not affect viability, but simultaneous deletion of *mgpB* and *corB* in the Δ*mgpA* background resulted in a drastic loss of viability comparable to that of the Δ*mgpA* Δ*corC* strain (Fig. 5D). Complementation with plasmid-borne *mgpB* or *corB* restored viability of the Δ*mgpA* Δ*mgpB* Δ*corB* strain (Fig. S5), confirming that loss of *mgpB* and *corB* underlies the stationary-phase lethality. Notably, all strains exhibiting severe loss of viability—Δ*mgpA* Δ*corC*, Δ*mgpA* Δ*mgpB* Δ*corB*, Δ*mgpA* Δ*mgpB* Δ*corC*, and Δ*mgpA* Δ*mgpB* Δ*corB* Δ*corC*—died to a similar extent (Fig. 5D), suggesting that these mutants fail in a common process required for successful transition into stationary phase.

**Fig. 5.**
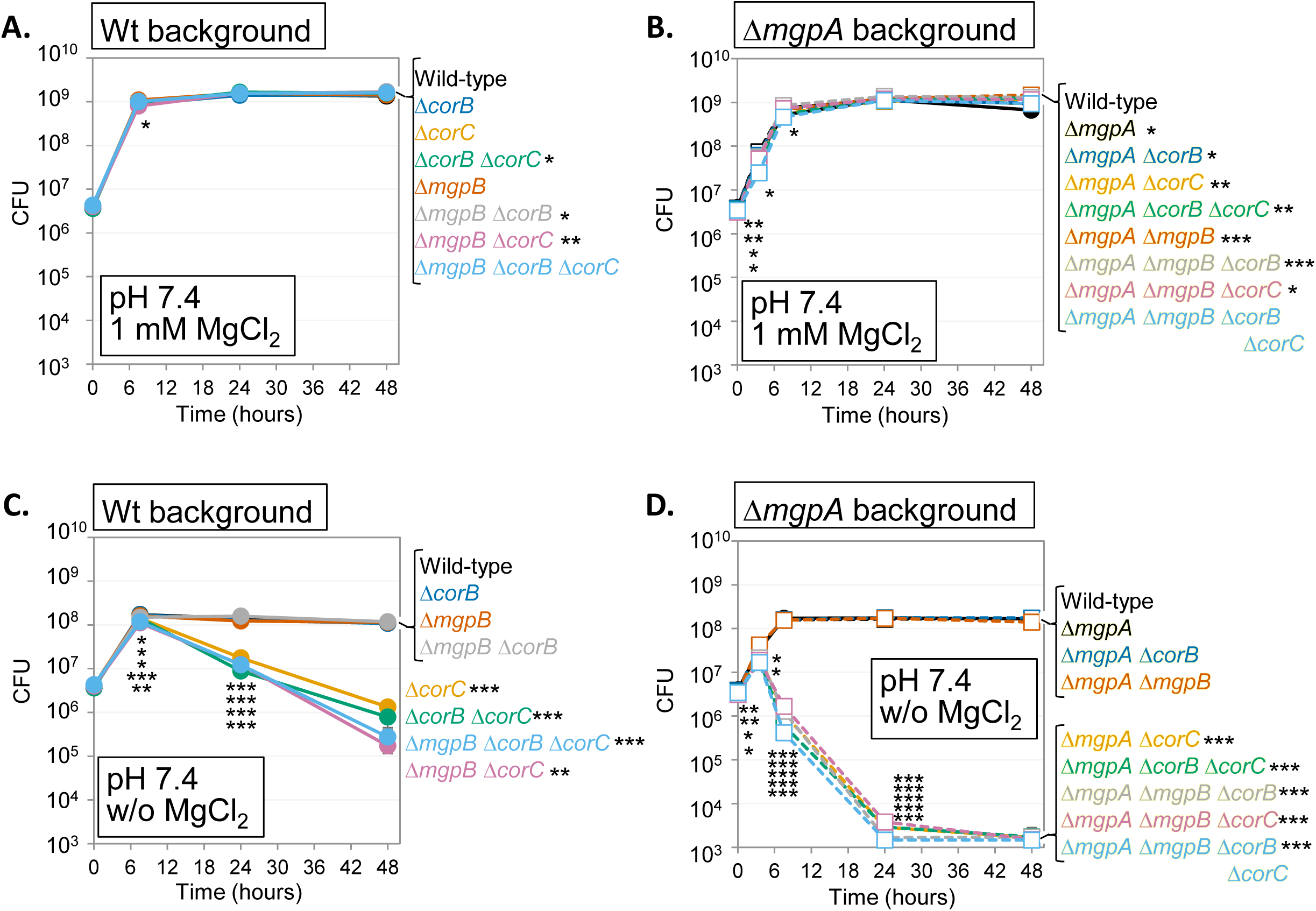
Simultaneous deletions of *mgpA*, *mgpB*, and *corB* cause loss of viability in stationary phase after Mg^2+^ starvation, similar to the double deletion of *mgpA* and *corC*. The indicated strains were grown overnight in N-minimal medium pH 7.4 with 1 mM MgCl_2_, washed, diluted into N-minimal medium pH 7.4 (A and B) with or (C and D) without 1 mM MgCl_2_ (t = 0 h), and incubated at 37℃. CFUs were determined at the indicated time points in (A and C) wild-type and (B and D) Δ*mgpA* background. Values are mean ± SD, n = 6. Unpaired t-test (P < 0.05*, 0.005**, 0.0005***) vs. wild type at the same time point and at the same Mg²^+^ concentration. Complete *p*-values for panels A-D are provided in Tables S6-9, respectively. Strains used: 14028, JS2740, JS2728, JS2741, JS2707, JS2742, JS2743, JS2744, JS2729, JS2745, JS2730, JS2746, JS2747, JS2737, JS2738, and JS2739.

To determine whether the loss of viability in the Δ*mgpA* Δ*mgpB* Δ*corB* strain is caused by excess-cation stress, we examined survival in backgrounds lacking either polyamine synthesis (Δsynth) or inducible high-affinity Mg²⁺ transport (Δ*mgtA* Δ*mgtB*), both of which suppress stationary-phase lethality in Δ*paeA*, Δ*corC,* and Δ*mgpA* Δ*corC* strains (15, 30). In the Δsynth background, the severe loss of viability of the Δ*mgpA* Δ*mgpB* Δ*corB* strain grown without added Mg²⁺ was partially suppressed (compare Fig. 5D and 6A), whereas in the Δ*mgtA* Δ*mgtB* background, this viability defect was completely suppressed (compare Fig. 5D and 6B). Deletion of *mgpA*, *mgpB*, and *corB* in the Δ*paeA* background further reduced stationary-phase viability (Fig. 6C), indicating PaeA and the MgpA/MgpB/CorB pathway contribute independently to the same physiological process. Together, these results demonstrate that MgpB and CorB are required to alleviate endogenous excess-cation stress that arises during stationary phase after Mg²⁺ starvation.

**Fig. 6.**
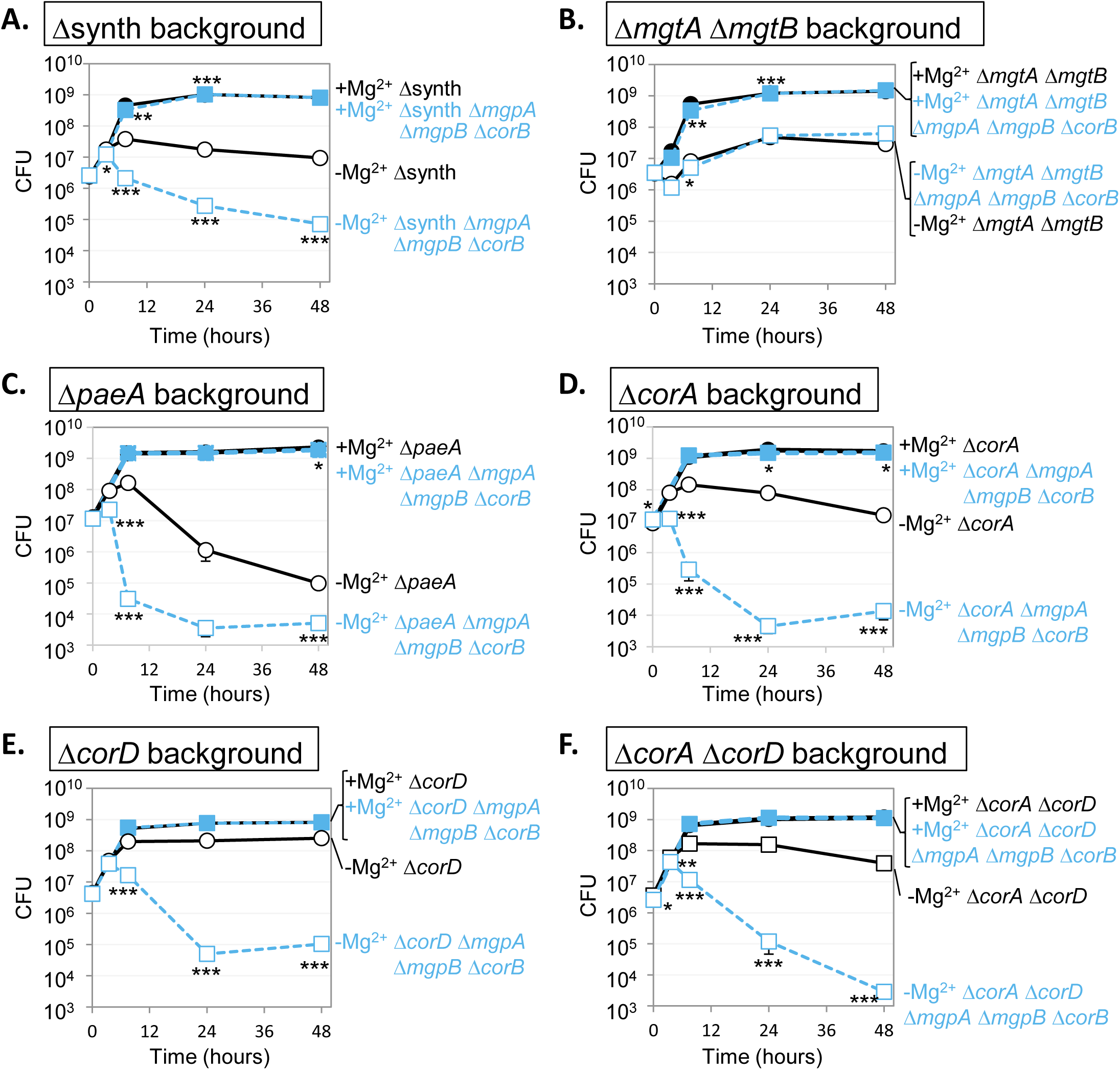
MgpA, MgpB, and CorB are required to alleviate endogenous excess-cation stress independently of PaeA, CorA, and CorD. The indicated strains were grown overnight in N-minimal medium pH 7.4 with 1 mM MgCl_2_, washed, diluted into N-minimal medium pH 7.4 (A and C) with or (B and D) without 1 mM MgCl_2_ (t = 0 h), and incubated at 37℃. CFUs were determined at the indicated time points in (A and B) wild-type and (C and D) Δ*mgpA* background. Values are mean ± SD, n = 6. Unpaired t-test (P < 0.05*, 0.005**, 0.0005***) vs. wild type at the same time point and at the same Mg²^+^ concentration. Complete *p*-values are provided in Table S14. Strains used: JS2560, JS2756, JS2713, JS2757, JS2720, JS2758, JS2695, JS2759, JS2760, JS2761, JS2762, and JS2763.

### CorA and CorD are dispensable for MgpA, MgpB, and CorB function

Previous work by Gibson et al. suggested that *corB* exhibits epistatic interactions with *corC*, *corD,* and *corA* (10). Indeed, the *corB* and *corC* genes exhibit an epistatic relationship during exogenous high-Mg^2+^ stress and endogenous excess-cation stress (Fig. 3C and 5D). To determine whether *corB* exhibits epistatic interactions with *corA* and *corD*, we examined the viability phenotype conferred by the Δ*mgpA* Δ*mgpB* Δ*corB* deletions during the transition to stationary phase after Mg²⁺ starvation in Δ*corA*, Δ*corD*, and Δ*corA* Δ*corD* backgrounds. CorA is the primary Mg²⁺ importer under Mg²⁺-replete conditions with the ability to mediate Mg²⁺ efflux under certain conditions (10). CorD (also known as ApaG) is a protein of unknown function found in both bacteria and eukaryotes (10, 40, 41).

In the Δ*corA* background, deletion of *mgpA*, *mgpB*, and *corB* caused a loss of viability comparable to that observed in the wild-type background (Fig. 6D), indicating that CorA is dispensable for the function of MgpA, MgpB, and CorB during endogenous excess-cation stress. This also shows that the CorA Mg^2+^ importer is not responsible for causing excess-cation stress. In the Δ*corD* background, deletion of *mgpA*, *mgpB*, and *corB* still resulted in a loss of viability upon entry into stationary phase after Mg²⁺ starvation (Fig. 6E), indicating that CorD is also dispensable for the essential function mediated by MgpA, MgpB, and CorB. However, the severity of the viability defect was partially suppressed compared to the wild-type background (compare Fig. 5D and Fig. 6E). This observation suggests that CorD contributes, either directly or indirectly, to the excess-cation stress that arises endogenously during the transition to stationary phase. The Δ*corA* Δ*corD* background exhibited a phenotype similar to that of the Δ*corD* single mutant (Fig. 6F), indicating that loss of *corA* does not further modify the effect of *corD* deletion. Collectively, these data show that CorA and CorD are dispensable for MgpA, MgpB, and CorB function.

### Deletions of *corB* and *mgpB* increase tolerance to exogenous polyamines

Previous work showed that PaeA, MgpA, and CorC all support survival during endogenous excess-cation stress in stationary phase after Mg²⁺ starvation (15, 30). PaeA is responsible for exporting divalent polyamines (33), whereas MgpA and CorC are responsible for reducing intracellular Mg²⁺ levels (15). Deletion of *paeA* sensitizes cells to exogenous polyamines (33), whereas deletion of *mgpA* or *corC* increases polyamine tolerance in the *paeA* background (15). Because the data presented above demonstrate that MgpB and CorB, like MgpA and CorC, are required for tolerance to exogenous Mg²⁺ and contribute to reducing cellular Mg²⁺ levels, we hypothesized that loss of these proteins might likewise enhance tolerance to exogenous polyamines. We therefore examined the effects of deleting *mgpA*, *mgpB*, *corB*, and *corC*, individually and in combination, on tolerance to exogenous polyamines. To increase the sensitivity of the assay, these mutations were introduced into a Δ*paeA* background, which is highly sensitive to polyamines (33), allowing additional effects on polyamine tolerance to be readily detected. Strains were grown to stationary phase in N-minimal medium (pH 7.4) containing low (50 μM), moderate (1 mM), or high (100 mM) Mg²⁺. These Mg²⁺ concentrations at neutral pH do not cause lethality. The cells were then washed and incubated in a pH 8.5 HEPES buffer supplemented with putrescine. Notably, alkaline pH promotes partial deprotonation of polyamines, facilitating their passive diffusion across the cytoplasmic membrane and into the cell (33, 42).

As expected, deletion of *paeA*, which encodes a divalent polyamine exporter (Fig. 7), sensitized cells to putrescine. Deletion of *mgpA*, *mgpB*, *corB*, and *corC*, either individually or in combination, generally increased polyamine tolerance, depending on the Mg^2+^ concentration in which the cells were grown (Fig. 7B and D). Consistent with our previous findings (15, 30), cells grown in moderate Mg²⁺ concentrations exhibit greater sensitivity to polyamines than cells grown with low Mg²⁺ (Fig. 7). Deletion of *mgpA* alone increased tolerance to putrescine only when cells were grown under low-Mg²⁺ conditions, whereas deletion of *mgpB* alone had no detectable effect on polyamine tolerance (Fig. 7B). In contrast, the Δ*mgpA* Δ*mgpB* double mutant exhibited increased tolerance to polyamines regardless of Mg²⁺ concentrations (Fig. 7B) and we used a higher putrescine concentration to address the effects of additional deletions in this background (Fig. 7D). Notably, this is the first time we observed a pronounced phenotype in the Δ*mgpA* Δ*mgpB* double mutant. These results indicate that MgpA and MgpB have a synergistic relationship. In the *mgpA*^+^ *mgpB*^+^ background, deletion of *corB* had little effect, whereas deletion of *corC* increased polyamine tolerance (Fig. 7B). In the Δ*mgpA* Δ*mgpB* background, deletion of *corB*, *corC*, or both genes increased polyamine tolerance to a similar extent (Fig. 7D). These results suggest that the functions of MgpA, MgpB, CorB, and CorC become detrimental when polyamines continuously enter the cell. The simplest explanation is that excess polyamines displace Mg²⁺ bound to macromolecules, increasing the cytoplasmic pool of free Mg²⁺. MgpA, MgpB, CorB, and CorC then promote Mg²⁺ efflux, leading to excessive loss of intracellular Mg²⁺ and ultimately cell death. Loss of Mg²⁺ efflux suppresses lethality, suggesting that Mg²⁺ and putrescine compete for binding. Indeed, growing the cells in high (100 mM) Mg²⁺ also partially suppresses lethality of certain strains, apparently providing addition competition against putrescine (Fig. 7).

**Fig. 7.**
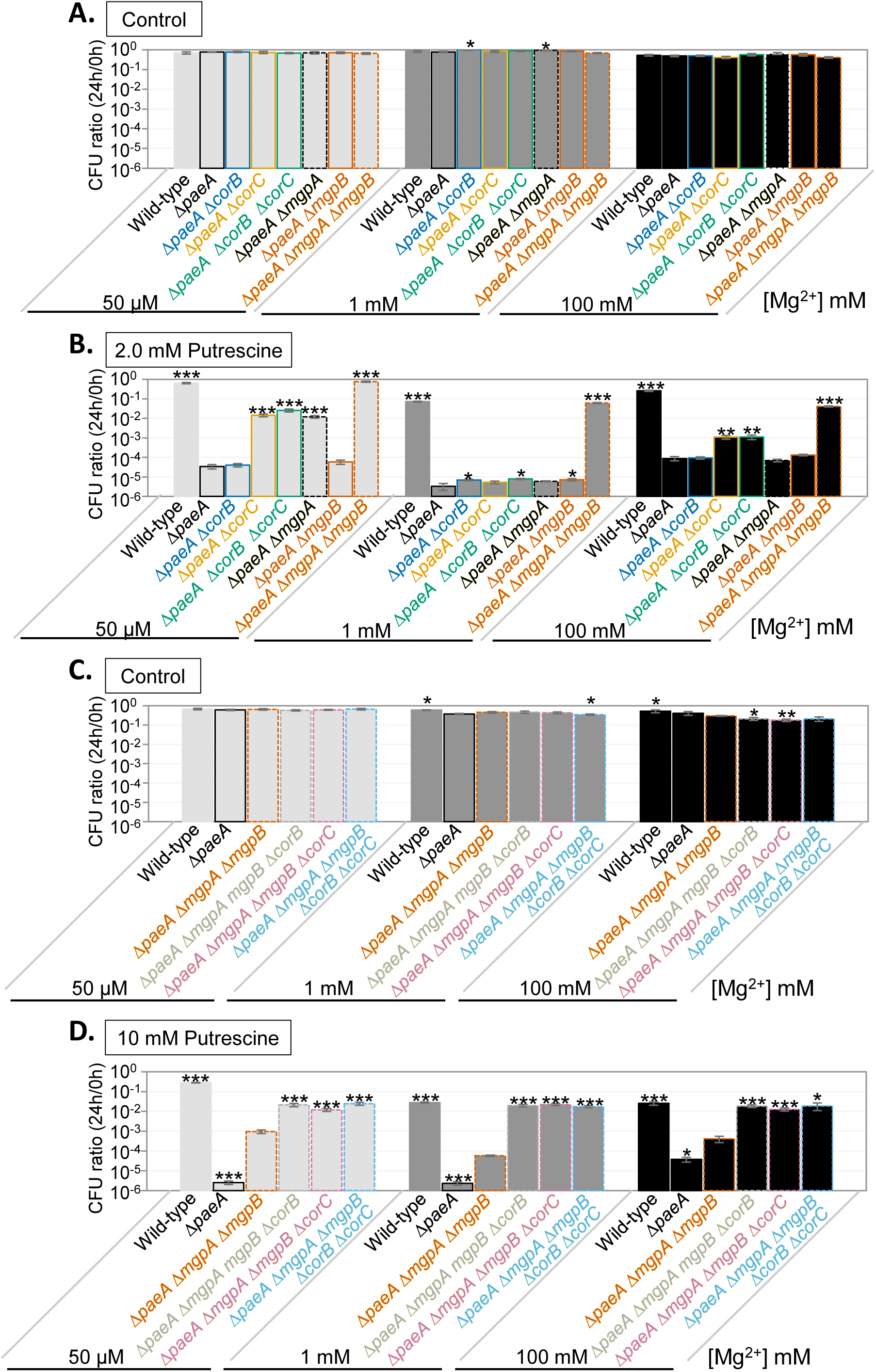
Deletions of *mgpA* and *mgpB* increase polyamine tolerance in Δ*paeA*, with further enhancement by *corB* deletion. The indicated strains (A and B) in Δ*paeA* (C and D) in Δ*paeA* Δ*mgpA* Δ*mgpB* backgrounds were grown in N-minimal medium (pH 7.4) with 50 μM, 1 mM or 100 mM MgCl_2_ for 24 hours at 37℃, washed, diluted into pH 8.5-buffered saline containing (A and C) no-polyamine control, (B) 2.0 mM putrescine, or (D) 10 mM putrescine (t=0h), and incubated at 37℃. CFUs were determined at 0h and 24h. CFU ratio (24h/0h) values are mean ± SD, n = 6. Unpaired t test (p < 0.05*, 0.005**, 0.0005***) versus (A and B) Δ*paeA*, or (C and D) Δ*paeA* Δ*mgpA* Δ*mgpB*. Strains used: JS2695, JS2779, JS2724, JS2780, JS2781, JS2759, JS2782, and JS2783.

### Effects of MgpA, MgpB, CorB, and CorC on Cobalt and Manganese Tolerance

Previously, we showed that deletion of *corC* increases tolerance to cobalt and manganese (15), consistent with earlier work identifying *corB*, *corC*, and *corD* as genes whose inactivation enhances cobalt tolerance (10). Despite the strong genetic interactions between *corC* and *mgpA*, deletion of *mgpA* alone does not increase cobalt and manganese tolerance (15). To further investigate the relationships among *mgpA*, *mgpB*, *corB*, and *corC* genes, we examined the effects of deletions of these deletions on cobalt and manganese tolerance. Deletion of *mgpA* alone had no effect on Mn²⁺ or Co²⁺ tolerance (Fig. S6), while deletion of *mgpB* alone conferred only a modest increase in Co²⁺ tolerance (Fig. S6). In contrast, simultaneous deletion of *mgpA* and *mgpB* increased tolerance to both metals, indicating that these proteins function redundantly. Deletion of *corC* increased Mn²⁺ and Co²⁺ tolerance regardless of the presence or absence of CorB, MgpA, or MgpB (Fig. S6), demonstrating that the effect of CorC is independent of these genes. In contrast, deletion of *corB* increased Mn²⁺ and Co²⁺ tolerance, especially in the Δ*mgpA* background. Together, these results indicate that MgpA and MgpB confer sensitivity to Co²⁺ and Mn²⁺, albeit to a lesser extent than CorB and CorC, which were discovered as by studying Co²⁺ tolerance (10).

### MgpB and CorB expression are regulated by Mg²⁺ levels

Our previous work showed that expression of both *mgpA* and *corC* is induced under high-Mg²⁺ conditions (15). Because *mgpB* and *corB*, like *mgpA* and *corC,* are required for tolerance to high-Mg²⁺ stress, we hypothesized that their expression is also regulated by Mg²⁺ availability. To test this hypothesis, we examined *mgpB* and *corB* expression using translational *lacZ* fusions. These constructs (Fig. 8A), which include the upstream regulatory regions and a portion of the coding sequence fused in frame to *lacZ*, were integrated at the λ *att* site on the chromosome, leaving the native *mgpB* and *corB* loci intact. Expression was measured after 24 h of growth in N-minimal medium (pH 7.4) containing various Mg²⁺ concentrations (0.05, 1, 10, or 100 mM MgCl₂) in wild-type and Δ*mgpA* Δ*mgpB* Δ*corB* backgrounds. Note that the Δ*mgpA* Δ*mgpB* Δ*corB* strain shows no growth or viability defects under these conditions (Fig. S7).

**Fig 8.**
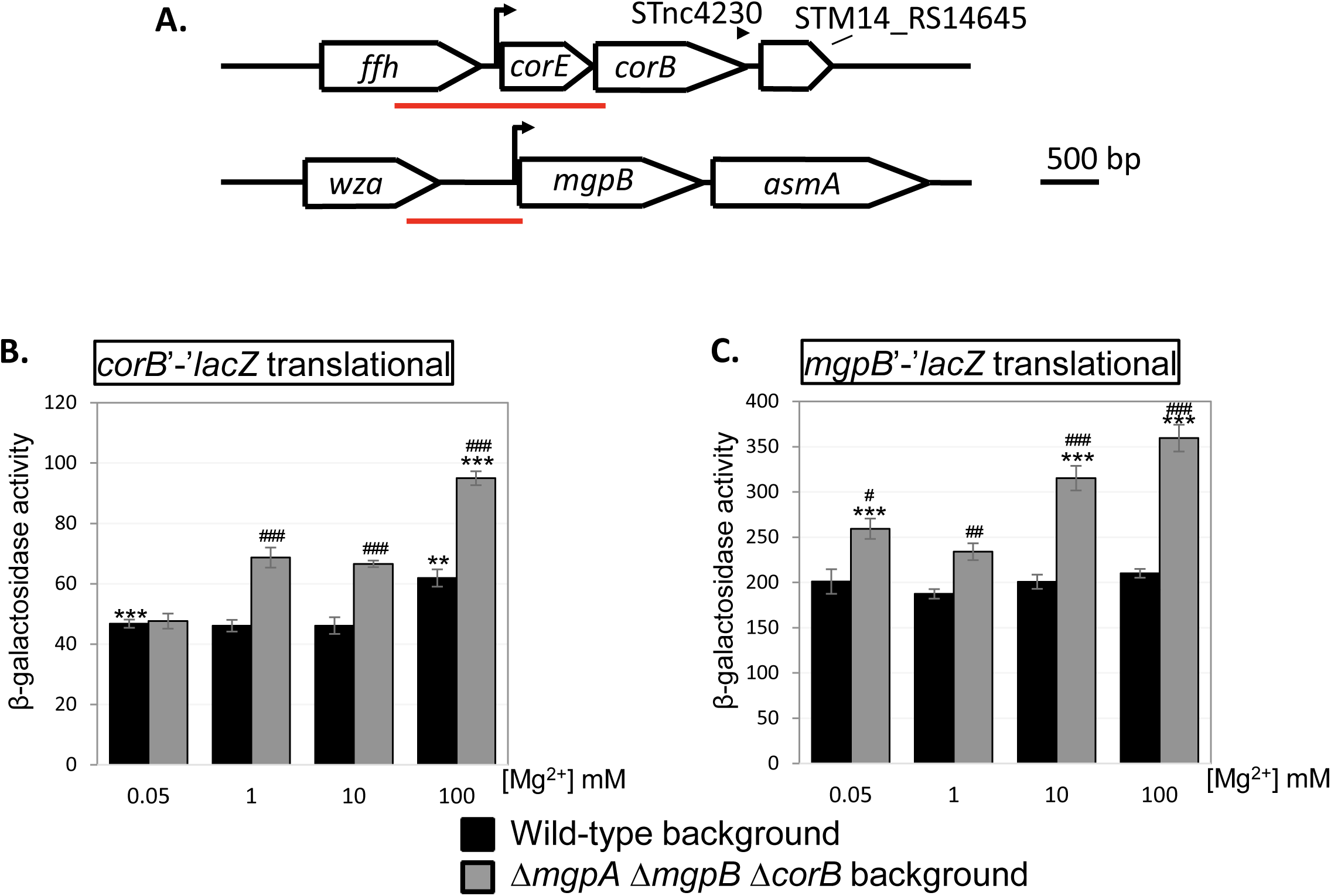
Expression of *corB* and *mgpB* are regulated by Mg^2+^ concentrations. (A) The *mgpB* and *corB* loci. Red bars indicate DNA fragments cloned upstream of *lacZ* for in-frame translational fusions. The (B) *corB-lac* translational fusion and (C) *mgpB-lac* translational fusion in the wild-type or Δ*mgpA* Δ*mgpB* Δ*corB* backgrounds were pre-grown to mid-exponential phase in N-minimal medium pH 7.4 with 1 mM MgCl_2_ at 37℃, washed, diluted in N-minimal medium pH 7.4 with or without the indicated amount of MgCl_2_, and incubated at 37℃ for 24h. β-galactosidase specific activity per mg protein was measured as described in Materials and Methods. Values are mean ± SD, n = 6. Unpaired t test (p < 0.05*, 0.005**, 0.0005***) versus 1 mM Mg^2+^ in the same background, (p < 0.05#, 0.005##, 0.0005###) vs wild-type background in the same condition. Strains used: JS2764, JS2765, JS2766, and JS2767.

In the wild-type background, the *corB* translational fusion showed an increase in expression when cells were grown with 100 mM MgCl₂ (Fig. 8B). To determine whether *corB* expression responds to elevated intracellular Mg²⁺, expression was measured in a Δ*mgpA* Δ*mgpB* Δ*corB* background, which accumulates substantially higher intracellular Mg²⁺ levels (Fig. 4B). In this background, expression of the *corB* translational fusion was higher than in the wild-type strain. Furthermore, expression increased 2-fold in 100 mM MgCl₂ relative to the low-Mg²⁺ condition (Fig. 8B). Together, these results suggest that elevated intracellular Mg²⁺ enhances *corB* expression through an as-yet-unknown mechanism.

The *mgpB* translational fusion in the wild-type background showed no increase in expression at higher extracellular Mg²⁺ concentrations (Fig. 8C). In contrast, in a Δ*mgpA* Δ*mgpB* Δ*corB* background, expression of *mgpB* increased in the presence of added Mg²⁺, reaching 1.2-fold and 1.7-fold higher levels in 1 mM and 100 mM MgCl₂, respectively, relative to in wild-type background. These results indicate that *mgpB* expression is upregulated when intracellular Mg^2+^ levels are elevated. Collectively, these data demonstrate that high intracellular Mg²⁺ upregulates MgpB and CorB expression through unknown mechanisms.

### MgpB and CorB are required for full tolerance to egg white

Our previous study demonstrated that the Δ*mgpA* Δ*corC* double mutant, but not either single mutant, is sensitive to egg white (15), an alkaline and viscous environment that contains multiple antimicrobial components (43). In contrast, in *S*. Enteritidis, deletion of *mgpA* alone is sufficient to confer egg white sensitivity (44). The specific egg white components responsible for killing the Δ*mgpA* Δ*corC* mutant remain unknown. However, high pH and Mg²⁺ likely contribute, as egg white is alkaline (pH 7.6 when fresh to ∼9.2 upon aging) and contains 3.7–4.9 mM Mg²⁺ (43, 45). Notably, the egg whites used in this study had a pH of approximately 8.6.

We hypothesized that MgpB and CorB also contribute to tolerance to egg white. To test this, we examined egg white tolerance in single, double, and triple mutants of *corB, mgpB*, and *mgpA* in both *corC*⁺ and Δ*corC* backgrounds. In the *corC*⁺ background, single deletion of *corB* or *mgpB*, double deletion of Δ*mgpB* and Δ*corB*, and double deletion of Δ*mgpA* and Δ*mgpB* double did not confer egg white sensitivity (Fig. 9A). In contrast, the Δ*mgpA* Δ*corB* mutant exhibited moderate sensitivity, whereas the Δ*mgpA* Δ*mgpB* Δ*corB* mutant showed severe sensitivity to egg white (Fig. 9A), mirroring their sensitivity to exogenous high-Mg²⁺ stress (Fig. 3C). In the Δ*corC* background, deletion of *corB*, *mgpB*, or both had no effect on survival, and all strains survived at levels comparable to the wild type. However, when *mgpA* was also deleted, all strains displayed similar severe sensitivity to egg white, equal to the *mgpA corC* strain (Fig. 9B). Complementation experiments confirmed these genetic requirements: introduction of plasmid-borne *mgpB* or c*orB* restored egg white tolerance in the Δ*mgpA* Δ*mgpB* Δ*corB* strain, while plasmid-borne *corB* conferred tolerance in the Δ*mgpA* Δ*corB* strain (Fig. S8). Collectively, these results demonstrate that MgpB and CorB function together with MgpA to promote tolerance to egg white, supporting a model in which multiple CorC-domain proteins cooperatively enhance egg white tolerance.

**Fig 9.**
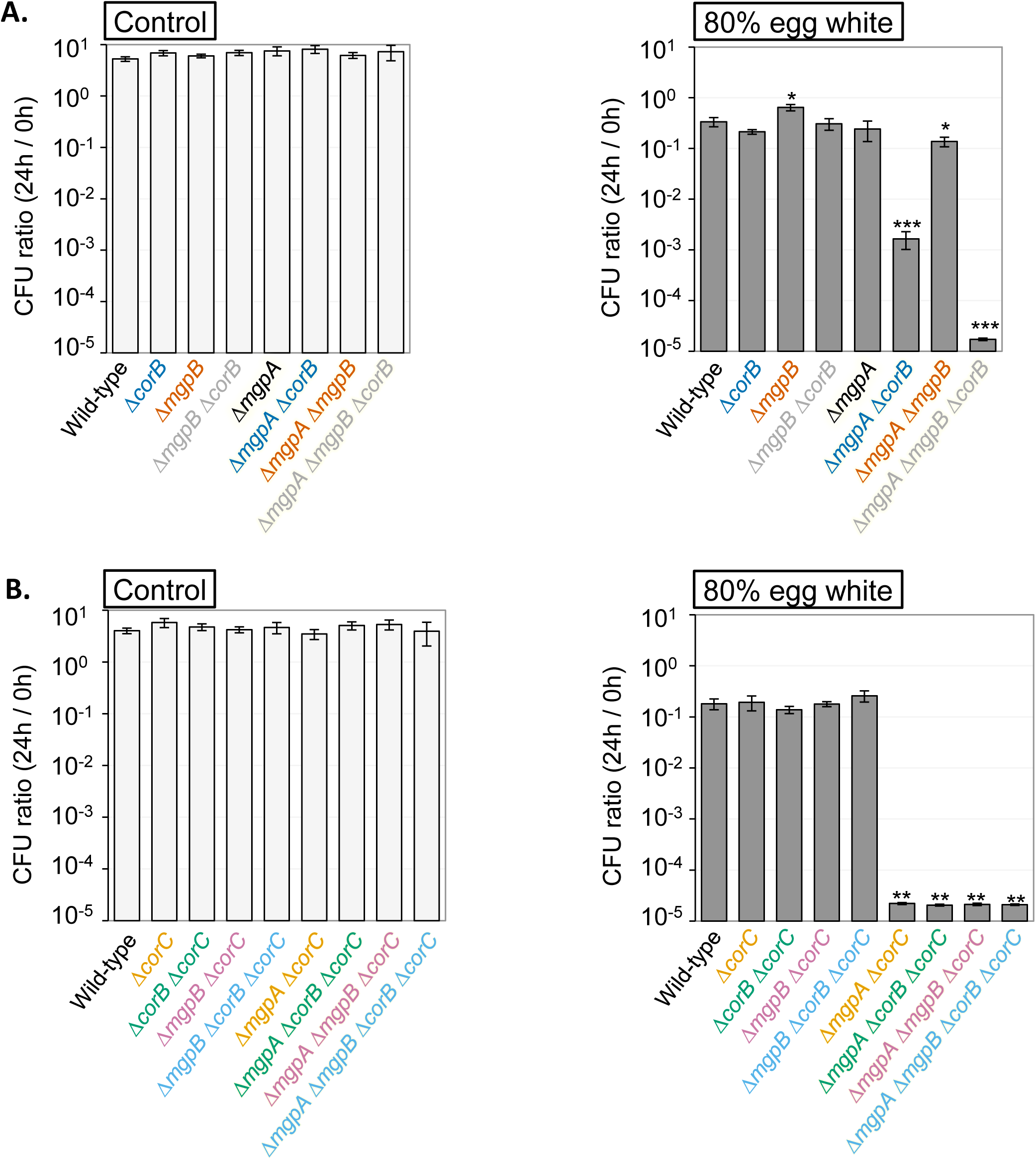
The Δ*mgpA* Δ*corB* strain is sensitive to egg white, and deletion of *mgpB* exacerbates the effect. The indicated strains were pre-grown to mid-exponential phase in N-minimal medium (pH 7.4) with 1 mM MgCl_2_ at 37°C, washed, diluted into (A and C) PBS buffered with 100 mM HEPES (pH 8.6) or (B and D) 80% egg white solution (t = 0 h), and incubated at 37°C. CFUs were determined at 0 hour and 24 hours. CFU ratio (24 h/0 h) values are mean ± SD, n = 6. Unpaired t-test (P < 0.05*, 0.005**, 0.0005***) versus WT same treatment. Strains used: 14028, JS2740, JS2728, JS2741, JS2707, JS2742, JS2743, JS2744, JS2729, JS2745, JS2730, JS2746, JS2747, JS2737, JS2738, and JS2739.

### MgpB and CorB are required for acute systemic infection

Successful *Salmonella* infection requires adaptation to the low-Mg²⁺ environment of the macrophage phagosome. This is underscored by the avirulence of a *phoP* mutant, which lacks the master regulator of the low-Mg²⁺ stress response (46, 47). We previously demonstrated that the CorC protein PaeA, a polyamine efflux transporter, as well as the other CorC proteins MgpA and CorC, which reduce cellular Mg²⁺ levels, are required for full virulence in mice (15, 30). These observations suggest that *Salmonella* experiences endogenous excess-cation stress and must reduce intracellular polyamines and Mg²⁺, similar to the response observed during stationary phase following in vitro Mg²⁺ starvation.

Therefore, we hypothesized that MgpB and CorB, which alleviate endogenous excess-cation stress by reducing cellular Mg^2+^ levels, also contribute to systemic infection. The contribution of MgpB and CorB to systemic infection was evaluated in BALB/c mice using competition assays. We focused on the Δ*mgpA* Δ*mgpB* Δ*corB* strain, which exhibits a severe loss of viability under endogenous excess-cation stress at stationary phase after Mg²⁺ starvation, conditions we assume that *Salmonella* encounters within macrophage phagosome. After intraperitoneal infection, the Δ*mgpA* Δ*mgpB* Δ*corB* strain exhibited attenuated virulence in BALB/c mice (Table 1), to a degree comparable to that of the Δ*mgpA* Δ*corC* strain (15). In contrast, the Δ*mgpA* single mutant was as virulent as the wild type (Table 1), indicating that *mgpA* inactivation alone does not explain the virulence defect. Instead, these results demonstrate that attenuation of the Δ*mgpA* Δ*mgpB* Δ*corB* and Δ*mgpA* Δ*corC* strains arises from the combined deletion of these genes, paralleling the observed genetic requirements for survival during stationary phase after Mg²⁺ starvation. Collectively, these findings identify MgpB and CorB as previously unrecognized contributors to acute systemic infection. Moreover, these findings suggest that the ability to reduce cellular Mg²⁺ levels is critical for *Salmonella* virulence.

**Table 1.** MgpB and CorB are important for the virulence of *Salmonella*.

| Genotype of<br>S. Typhimurium strain<sup>a</sup> |  | Organ | Median CI<sup>b</sup> | No. of mice | p value<sup>c</sup> | Fold<br>change<sup>d</sup> |
| --- | --- | --- | --- | --- | --- | --- |
| *ΔmgpA ΔmgpB ΔcorB* | WT | Spleen | 0.07 | 10 | <0.0001 | 12.9 |
|  |  | Liver | 0.10 | 10 | <0.0001 | 9.2 |
| *ΔmgpA* | WT | Spleen | 2.14 | 5 | NS | 0.47 |
|  |  | Liver | 2.21 | 5 | NS | 0.46 |

<sup>a</sup>Strains used were 14028, JS2737, and JS2693.
<sup>b</sup>Bacteria were recovered from the spleen and liver after intraperitoneal (i.p.) competition assays. The competitive index (CI) was calculated as (percent strain A recovered/percent strain B recovered)/(percent strain A inoculated/percent strain B inoculated).
<sup>c</sup>A nonparametric test, the Mann–Whitney U test, was used to compare the CI values to the inocula. NS, not significant.
<sup>d</sup>The fold attenuation is 1 /CI, reflecting the level of attenuation of strain A relative to strain B.

## DISCUSSION

Upon Mg²⁺ starvation, *Salmonella* induces both polyamine synthesis and high-affinity Mg²⁺ transport (Fig. 1) (30, 31). Either pathway alone is sufficient to support survival, whereas simultaneous disruption of both leads to cell death. However, upon entry into stationary phase, the combination of polyamine synthesis and high-affinity Mg²⁺ transport leads to endogenous excess-cation stress (Fig. 1). If the cell cannot alleviate this stress, viability is lost through mechanisms that remain unclear (30). CorC domain-containing proteins play a central role in alleviating this stress. We previously showed that PaeA mitigates excess-cation stress by exporting the divalent polyamines putrescine and cadaverine (30), and that MgpA (YoaE) and CorC relieve excess-cation stress by reducing cellular Mg²⁺ (15). In this study, we identify MgpB (YegH) and CorB as acting redundantly and synergistically with MgpA and CorC to maintain cation homeostasis by reducing Mg^2+^ levels in stationary phase. These four proteins act distinctly from PaeA, which is responsible for reducing cellular polyamine levels (30). MgpA, MgpB, CorB, and CorC, but not PaeA, also confer tolerance to egg white (Fig. 9) (15), an environment that facilitates *Salmonella* transmission between hosts, suggesting that disruption of Mg²⁺ homeostasis is one mechanism underlying the antimicrobial activity of egg white. Mg²⁺ limitation is a hallmark of the macrophage phagosome, yet, like Mg²⁺ starvation *in vitro*, endogenous excess-cation stress appears to occur in *Salmonella* during replication in macrophages. Thus, MgpB and CorB, similar to the other CorC-domain-containing proteins PaeA, MgpA, and CorC (15, 30), play a role during acute systemic infection (Table 1). Together, our findings establish a framework for understanding excess-cation stress and identify MgpB and CorB, in addition to MgpA and CorC, as proteins required for reducing cellular Mg²⁺ levels, a process that has remained poorly understood in bacteria.

Loss of MgpA, MgpB, CorB, and CorC results in increased intracellular levels of Mg²⁺ and the synergistic phenotypes suggest that they act redundantly to export excess Mg^2+^ from the cell. MgpA and CorC seem to play the dominant roles, with subtle differences is importance depending on the various phenotypes that we examined. Loss of MgpB and/or CorB often had significant phenotypes only in the absence of MgpA in particular. In *B. subtilis* and *S. aureus*, deletion of *corB* homologs causes Mg²⁺ accumulation and high-Mg²⁺ sensitivity, whereas *Salmonella corB* mutations alone confer neither phenotype. This difference likely reflects the presence of MgpA and MgpB in *Salmonella*, which are absent from these other organisms. We note that CorC does not have a transmembrane domain, suggesting that it regulates some other transmembrane protein to control Mg^2+^ efflux. The fact that *corC* mutations are not simply epistatic with loss of any of the other CorC-domain proteins (15) strongly suggests that the putative CorC partner is some additional unknown protein.

Cells grown in low Mg²⁺ were more resistant to polyamines in stationary phase than cells grown 1 mM Mg²⁺ (Fig. 7), suggesting that when Mg²⁺ is limited, polyamines are less toxic, consistent with our previous report (30). However, it is also clear that cation stress is not simply the result of higher combined concentrations of Mg^2+^ and polyamines. Rather, it is the proper coordination among intracellular cations that is essential for maintaining cellular viability. For example, elevated intracellular K⁺ increases tolerance to excess polyamines (48). In addition, K⁺ influx and Mg²⁺ efflux occur simultaneously under osmotic stress (49), suggesting coordinated regulation of, and competition between, these cations. Cation competition also explains why loss of the putative Mg^2+^ exporters suppresses the sensitivity of a *paeA* mutant to polyamines.

During polyamine treatment under alkaline conditions, polyamines continue to enter the cytoplasm. The elevated intracellular polyamine concentrations apparently compete with Mg²⁺ for binding to negatively charged macromolecules, thereby displacing Mg²⁺ from these cellular reservoirs. The released Mg²⁺ is then exported by MgpA, MgpB, CorB, CorC, leading to a reduction in Mg²⁺ and a further imbalance between Mg^2+^ and polyamines. Loss of the Mg^2+^exporters keeps the Mg^2+^ high, allowing it to compete with the polyamines. This implies that it is the binding of polyamines, rather than Mg^2+^, to some particular anionic compound or subset of compounds that is lethal, rather than some overall increase in cation. Consistent with this, cells grown in 100 mM Mg²⁺ were actually more tolerant to polyamines than those grown with 1 mM Mg²⁺ (Fig. 7). Similarly, when Mn²⁺ or Co²⁺ enters the cytoplasm, primarily through CorA-dependent pathways (50, 51), these excess cations may compete with Mg²⁺ for intracellular binding sites, leading to displacement of Mg²⁺ from cellular ligands. Consistent with the Irving– Williams series, the thermodynamic stability of complexes formed between these metals and biological ligands generally increases in the order Mg²⁺ < Mn²⁺ < Fe²⁺ < Co²⁺ (52). Under normal physiological conditions, the Mg^2+^ concentration is maintained at a concentration at least an order of magnitude above the concentration of these other cations (53, 54). In response to excess Mn²⁺ or Co²⁺, CorB, CorC, MgpA, and MgpB act to restore intracellular cation homeostasis by promoting Mg²⁺ export, which exacerbates Mg²⁺ depletion, resulting in reduced cell viability (Fig. S6). A similar phenomenon has been reported in *B. subtilis*, where CorB homolog MpfA-mediated Mg²⁺ efflux contributes to Mn^2+^ and Co²⁺ toxicity (13). Note, however, that the relative role of the putative Mg²⁺ exporters differs in response to Mn^2+^ or Co²⁺, with loss of CorC and CorB having a much greater effect than loss of MgpB and MgpA. This implies that there might be differential regulatory effects on either expression or function of the various exporters in response to Mn^2+^ or Co²⁺, perhaps explaining the need for multiple, otherwise redundant, exporters.

Our data suggest that MgpA, MgpB, CorB, and CorC are acting redundantly to export Mg²⁺ from the cell. However, given the fact that it is the competition between cations that is critical to overall viability, we cannot rule out the possibility that these exporters contribute to reducing cellular Mg²⁺ levels indirectly by transporting other cations or anions that influence Mg²⁺ homeostasis. Our data, together with the homology of CorB to a characterized Mg²⁺ efflux transporter (11, 12), support a model in which CorB functions directly as an Mg²⁺ efflux transporter. In contrast, the TerC transmembrane domain, which is present in MgpA and MgpB, has been implicated in Mn²⁺ transport in *B. subtilis* (39, 55). Increased intracellular phosphate levels can also reduce free Mg²⁺ by promoting ATP synthesis (56), which requires Mg²⁺ for charge neutralization. In fact, *mgpB* in *E. coli* is regulated by PhoB, a master regulator of phosphate starvation (57). Further biochemical analyses are required to prove that MgpA and MgpB transport Mg²⁺ per se.

All CorC-domain proteins in *Salmonella* contribute to maintaining cation homeostasis. Despite these physiological roles, the molecular function of the conserved CorC domain remains unknown. The tandem CBS domains located upstream of the CorC domain bind and coordinate Mg^2+^-ATP at the dimer interface without hydrolyzing it, thereby promoting dimerization and activation of transporter activity (12, 36). Thus, we hypothesize that the CBS pair domains sense intracellular Mg²⁺ availability through Mg²⁺-ATP binding. We speculate that the CorC domain may provide an additional regulation beyond, or in combination with, Mg^2+^-ATP-dependent activation.

Expression of MgpB and CorB is upregulated under conditions of elevated Mg²⁺ (Fig. 8B and C), providing additional support for the idea that these proteins protect cells from high-Mg²⁺ stress. Deletion of *mgpA*, *mgpB*, and *corB* exaggerates the effect of high Mg²⁺ on the expression of MgpB and CorB. These mutations increase the accumulation of Mg²⁺ in cells (Fig. 4B), suggesting that regulation of CorB and MgpB responds primarily to intracellular Mg²⁺ levels rather than extracellular Mg²⁺ concentrations. Comprehensive RNA-seq analyses in *Salmonella* indicate that *corB* is upregulated ∼3.6-fold and *mgpB* ∼1.4-fold within macrophages (58), consistent with the concept that *Salmonella* undergoes excess-Mg^2+^ stress during infection. Transcriptomic analyses further show that neither gene is affected by PhoPQ regulation; however, *mgpB* is downregulated ∼1.5-fold in the deletion mutant of *slyA* (26), which is a transcriptional regulator required for survival within macrophages (59). Together, these findings suggest that *corB* and *mgpB* respond to high-Mg²⁺ conditions, although the underlying regulatory mechanisms that integrate Mg²⁺ signals remain unclear and require further investigation.

Disruption of Mg^2+^ homeostasis ultimately causes cell death (Fig. 3C and 5D) by mechanisms that remain to be elucidated. CorC has been implicated in ribosome homeostasis (60, 61), and homologs of CorB have been linked to ribosome homeostasis and RNA homeostasis (62, 63). In *E. coli*, a Δ*corC* mutant accumulates aberrant 16S rRNA at stationary phase through a unknown mechanism (60). In *B. subtilis*, inactivation of the CorB homolog MpfA suppresses the growth defect of a ΔL34 ribosomal protein mutant (63). In *S. aureus*, inactivation of the CorB homolog MpfA suppresses the slow-growth phenotype of a mutant of *cshB*, which encodes a DEAD-box RNA helicase required for post-transcriptional responses to cold shock (14, 62, 64). Our findings suggest that these effects are indirect and that excess-Mg²⁺ stress impairs ribosome integrity and RNA function. Further study is required to understand the toxicity of excess-Mg^2+^ stress.

## EXPERIMENTAL PROCEDURES

### Media

For routine strain construction, bacteria were cultured at 37°C in Luria-Bertani (LB) medium containing 1% NaCl. Strains harboring the temperature-sensitive plasmids pCP20 or pKD46 were maintained at 30°C. Antibiotics were added when appropriate at the following final concentrations: ampicillin, 50 μg/ml; chloramphenicol, 10 μg/ml; kanamycin, 50 μg/ml; and apramycin, 50 μg/ml. N-minimal medium was prepared as described previously (23, 65) and contained 5 mM KCl, 7.5 mM (NH_4_)_2_SO_4_, 0.5 mM K_2_SO_4_, 1 mM KH_2_PO_4_, 0.1 M Tris, 15 mM glycerol, and 0.1% casamino acids. The pH and Mg^2+^ concentrations were adjusted as indicated. MOPS medium was prepared as described previously (66) and contained 40 mM MOPS, 4 mM Tricine, 9.5 mM NH₄Cl, 0.276 mM K₂SO₄, and 50 mM NaCl. The medium was adjusted to pH 7.4 with KOH and supplemented with 15 mM glycerol and 0.1% casamino acids. When indicated, 1.32 mM K₂HPO₄ and 10 mM MgCl₂ were added. Trace metals from the original formulation were omitted to minimize the effects of metal supplementation.

### Bacterial strains and plasmids

The bacterial strains and plasmids used in this study are listed in Tables S1 and S2, respectively. All *Salmonella* strains were derived from *Salmonella enterica* serovar Typhimurium 14028. Gene deletions with insertion of antibiotic resistance cassettes were generated by λ Red-mediated recombination as described previously (67, 68), with the endpoints specified in Table S1. Deletions were verified by PCR and transduced into the desired strains by P22 HT105/1 int-201-mediated transduction (69). All plasmids were passaged through the restriction-minus, modification-plus Pi+ strain JS198 (68) before transformation into *Salmonella*. Antibiotic resistance markers were excised using pCP20, and the resulting mutants were verified by PCR.

For construction of the translational *corB’-‘lacZ* and *mgpB’-‘lacZ* fusions, the corresponding regulatory regions, including the third and first amino acids of the *corB* and *mgpB* open reading frames, respectively, were amplified by PCR with the primers listed in Table S3. The PCR products were fused upstream and in frame with the promoterless *lacZ* gene in pDX1 vector using NEBuilder® HiFi DNA Assembly (Gibson assembly) (Fig. 8). The resulting products were introduced into DH5α λ*pir*, and their sequences were verified by DNA sequencing. The pDX1-*corB*-*lacZ* and pDX1-*mgpB*-*lacZ* were subsequently integrated into the λ attachment site of the *Salmonella* chromosome as described previously (69). To construct pWKS30-p*lacZ*-*corB* and pWKS30-p*lacZ*-*mgpB*, the *corB* and *mgpB* regions were individually amplified by PCR and assembled with PCR-amplified pWKS30 using NEBuilder® HiFi DNA Assembly. The primers used are provided in Table S3. The resulting products were transformed into DH5α λ*pir* and their sequences were verified by DNA sequencing. The cloned regions and their endpoints are listed in Tables S2 and S3.

### Mg^2+^ sensitivity assay

Mg²⁺ sensitivity was measured as described previously (15). Overnight cultures grown in N-minimal medium (pH 7.4) supplemented with 1 mM MgCl₂, were diluted 800-fold into the same medium and grown for 4.5 hours. Cells were harvested, washed three times with PBS, and adjusted to an OD_600_ of 0.005 in N-minimal medium (pH 8.5) containing the indicated concentrations of MgCl_2_. After incubation at 37°C in 13 x 100-mm test tubes, the OD₆₀₀ of the culture was measured using a BioTek ELx808 Absorbance Reader at the indicated time points. When necessary, cultures were diluted in the original growth medium to ensure accurate OD₆₀₀ measurements. To determine CFU, serial dilutions of the cultures were plated on LB agar plates and incubated overnight at 37°C.

### Bioassay for Mg^2+^

As described previously, intracellular Mg²⁺ levels were measured by bioassay (15). To prepare the lysates, overnight cultures grown in N-minimal medium (pH 7.4) supplemented with 1 mM MgCl₂ were diluted 800-fold into the same medium and grown for 4.5 hours. The cells were then washed three times with PBS, adjusted to an OD₆₀₀ of 0.005 in 15 mL of N-minimal medium (pH 7.4) containing 10 mM MgCl₂. Cultures were incubated at 37°C in 125-mL baffled flasks for 24 h. To minimize cation efflux, chloramphenicol was added to 10 μg/ml for 5 min before harvest, after which the cultures were placed in ice-cold water for an additional 5 min. Cells were collected at 7,000 × g, washed once with PBS containing 100 mM EDTA, and washed twice more with Chelex-treated PBS. The resulting pellets were resuspended in 350 μL Chelex-treated PBS and disrupted by sonication. Debris was removed by centrifugation (12,000 × g). Protein concentrations in the clarified lysates were determined using the Bradford assay (Bio-Rad) and adjusted to 125 mg/ml with Chelex-treated PBS. Samples were subsequently heated at 98°C for 15 min to denature proteins and release Mg^2+^.

To assess the growth-stimulating activity of the cell lysates, overnight cultures grown in N-minimal medium (pH 7.4) supplemented with 10 mM MgCl₂ were diluted 800-fold into the same medium and grown for 4.5 hours. Cells were washed three times with PBS and adjusted to an OD_600_ of 0.005 in 250 μL of N-minimal medium. Cultures were supplemented either with the indicated concentrations of MgCl_2_ or with 34 μg/ml of the cell lysate in the presence or absence of 1 mM MgCl_2_. Cultures were incubated in a BioTek ELx808 Absorbance Reader with shaking for 15 hours, after which final OD_600_ was measured. When necessary, cultures were diluted in the original growth medium to ensure accurate OD₆₀₀ measurements. Cellular Mg^2+^ concentrations in the test cells were calculated from the OD₆₀₀ measured at 15 h using the equation derived from the calibration curves, assuming 1.55 × 10⁻¹⁵ g of protein per cell (BioLabs) and a cell volume of 2.3 × 10⁻¹⁵ L per cell (70).

### Survival assay

Overnight cultures in N-minimal medium (pH 7.4) containing 1 mM MgCl_2_ were diluted 1:800 into the same medium and incubated for 4.5 h. After three washes with PBS, the cultures were adjusted to an OD_600_ of 0.005 in 1 mL of N-minimal medium at pH 7.4 containing either no added MgCl_2_ or 1 mM MgCl_2_, and incubated at 37°C in 13 × 100-mm test tubes. At the indicated time points, aliquots were serially diluted and plated on LB agar. Plates were incubated overnight at 37°C, and CFU were determined. For growth measurements, OD_600_ values were recorded using a BioTek ELx808 Absorbance Reader.

### Mn^2+^ and Co^2+^ sensitivity assays

Overnight cultures in modified MOPS medium (pH 7.4) supplemented with 1.32 mM K_2_HPO_4_ and 1 mM MgCl_2_ were washed three times with PBS and adjusted to an OD_600_ of 0.05 in 200 μL of modified MOPS medium containing 15 mM glycerol. When indicated, MnCl_2_ or CoCl_2_ was added to final concentrations of 1 mM or 0.2 mM, respectively. Cultures were incubated in 96-well microplates at 37°C with shaking for 9 h. To determine CFU, before and after incubation, serial dilutions of the cultures were spotted onto LB agar plates, which were subsequently incubated overnight at 37°C.

### Polyamine sensitivity assay

Putrescine stock solution was prepared at 500 mM in 0.85% NaCl buffered to pH 8.5 with HEPES. Overnight cultures grown in N-minimal medium (pH 7.4) containing 10 mM MgCl_2_ were washed three times with 0.85% NaCl, adjusted to an OD_600_ of 0.005 in 15 ml of N-minimal medium containing 50 μM, 1 mM, or 100 mM MgCl_2_, and incubated at 37°C for 24 h in 125-ml baffled flasks. Cells were washed three times and resuspended in an equivalent volume of 0.85% NaCl containing 100 mM HEPES (pH 8.5). Aliquots of 200 μL were transferred to 96-well microplates, and putrescine was added to the indicated final concentration. Plates were incubated at 37°C with shaking at 200 rpm. Before and after incubation, serial dilutions of the cultures were spotted onto LB agar plates, which were subsequently incubated overnight at 37°C to determine CFU.

### β-galactosidase assay

Overnight cultures in N-minimal medium (pH 7.4) containing 1 mM MgCl₂ were diluted 800-fold into the same medium and grown for 4.5 h. Cells were washed three times with PBS and adjusted to an OD₆₀₀ of 0.005 into 15 ml of N-minimal medium (pH 7.4) containing the indicated concentration of MgCl₂. Cultures were then incubated at 37°C in 125-ml baffled flasks for 24 h. Chloramphenicol at the final concentration of 10 μg/mL was added to the culture and incubated for 5 minutes, followed by transfer of the flasks to ice-cold water for another 5 minutes. Then, cells were collected by centrifugation (7,000 × g), resuspended in Z buffer, and disrupted by sonication. Cell debris was removed by centrifugation (12,000 × g), and the resulting lysates were assayed for β-galactosidase activity in a microtiter plate format, as previously described (71). Total protein concentration was measured by a Bradford assay (Bio-Rad). The β-galactosidase activity units are defined as (μmol of ortho-nitrophenol formed per minute) / mg protein and are presented as the mean ± standard deviation with n = 6.

### Egg white sensitivity assay

Tolerance to egg white was measured as described previously (15). Eggs were obtained from the Meat & Egg Sales Room of the Department of Animal Sciences at the University of Illinois at Urbana-Champaign and stored at 4°C for less than 1 month before use. Egg white was collected from the eggs and mixed with a 1:4 volume of 0.85% NaCl. The mixture was homogenized on ice using a mortar and pestle, followed by centrifugation to remove debris. The resulting solution was used as an 80% egg white solution. Note that these egg white solutions did not contain microbes capable of growing on an LB plate.

Overnight cultures were diluted 800-fold into N-minimal medium (pH 7.4) containing 1 mM MgCl_2_ and grown for 4.5 h. Cells were washed three times with 0.85% NaCl and adjusted to an OD_600_ of 0.02 in 1 ml of either 80% egg white solution or 0.85% NaCl containing 100 mM HEPES (pH 8.5) in 13 x 100-mm test tubes. The cultures were incubated at 37°C on a roller drum. At 0 and 24 h, serial dilutions of the cultures were plated on LB agar plates, which were incubated overnight at 37°C to determine the CFU.

### Animal assay

All animal experiments were reviewed and approved by the University of Illinois Institutional Animal Care and Use Committee (IACUC) and conducted under protocol no. 24171. The competition experiments were performed using 5-to 6-week-old BALB/cAnNHsd mice obtained from Envigo. *Salmonella* strains were grown separately overnight in LB medium, mixed at a 1:1 ratio, and diluted in sterile PBS to obtain a target inoculum of approximately 400 CFU in a 200-μL volume. Mice were infected intraperitoneally. An aliquot of each inoculum was plated on LB agar to determine the total bacterial load, and colonies were replica-plated onto the appropriate selective media to calculate the input ratio for each strain. Four days after infection, BALB/cAnNHsd mice were euthanized by CO_2_ asphyxiation followed by cervical dislocation. Spleens and livers were collected and homogenized. Serial dilutions of the tissue homogenates were plated on LB agar and incubated overnight. The resulting colonies were replica-plated onto the appropriate selective medium to calculate the output ratio for each competition. The competitive index (CI) was calculated as (percent strain A recovered/percent strain B recovered)/(percent strain A inoculated/percent strain B inoculated). The statistical comparisons of individual competitions were performed using a nonparametric Mann-Whitney test.

## ACKNOWLEDGEMENTS

We are grateful to Yekaterina Golubeva for critically reading the manuscript. This study was supported by grant R21 AI199588 to JMS.

