## Supplemental Materials for "CorB and MgpB (YegH) alleviate excess-magnesium stress and contribute to egg white tolerance and virulence in *Salmonella*"

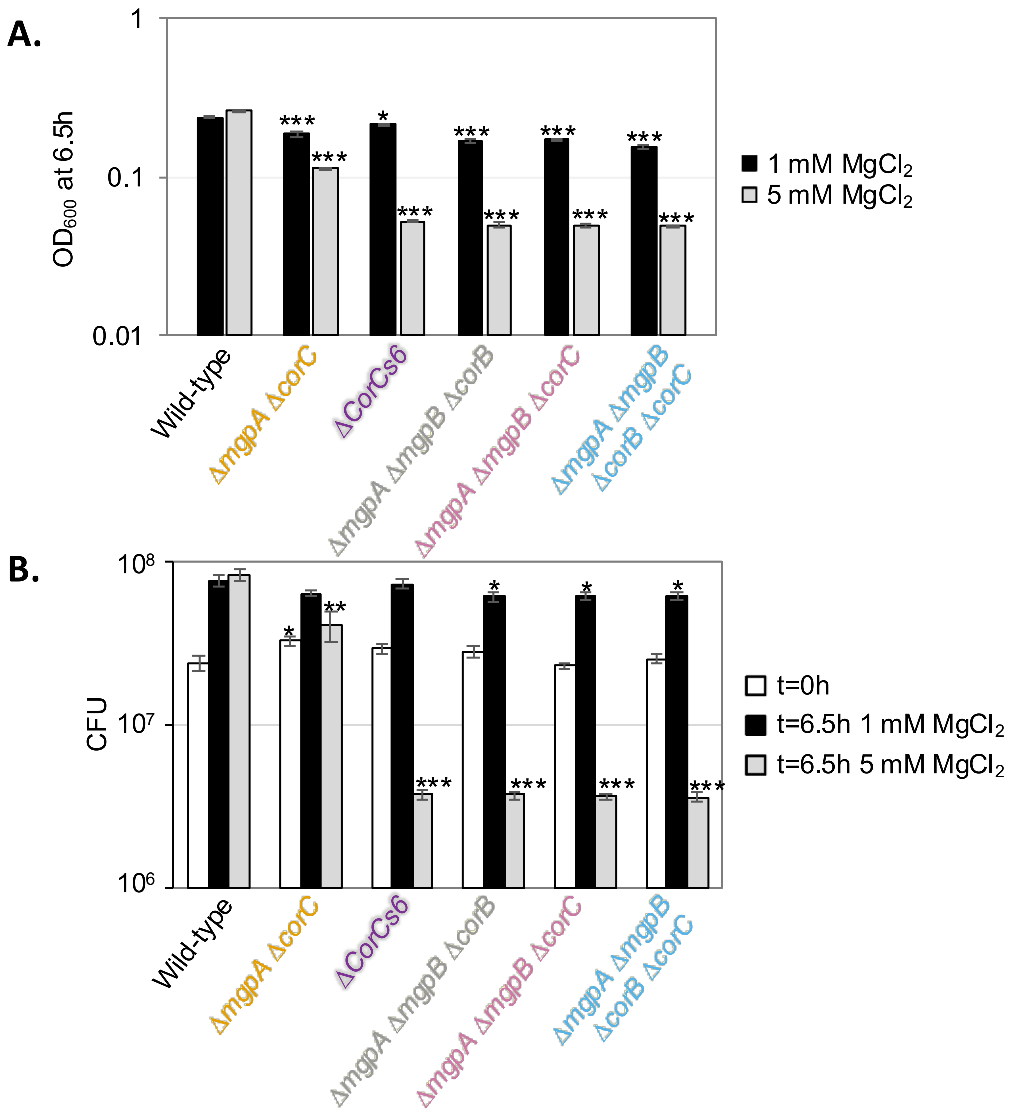


**Fig. S1. Loss of MgpB or CorB sensitizes cells to exogenous high $Mg^{2+}$ at high pH in the absence of MgpA.** The indicated strains were pre-grown to mid-exponential phase in N-minimal medium pH 7.4 with 1 mM $MgCl_2$, washed, and diluted into N-minimal medium pH 8.5 with 1 or 5 mM of $MgCl_2$, followed by incubation at 37 ℃. (A) $OD_{600}$ and (B) CFU were measured after 6.5 hours of incubation. Values are mean ± SD, n = 6. Unpaired t test (p < 0.05*, 0.005**, 0.0005***) versus corresponding WT at the same condition. Strains used: 14028, JS2730, JS2736, JS2737, JS2738, and JS2739.

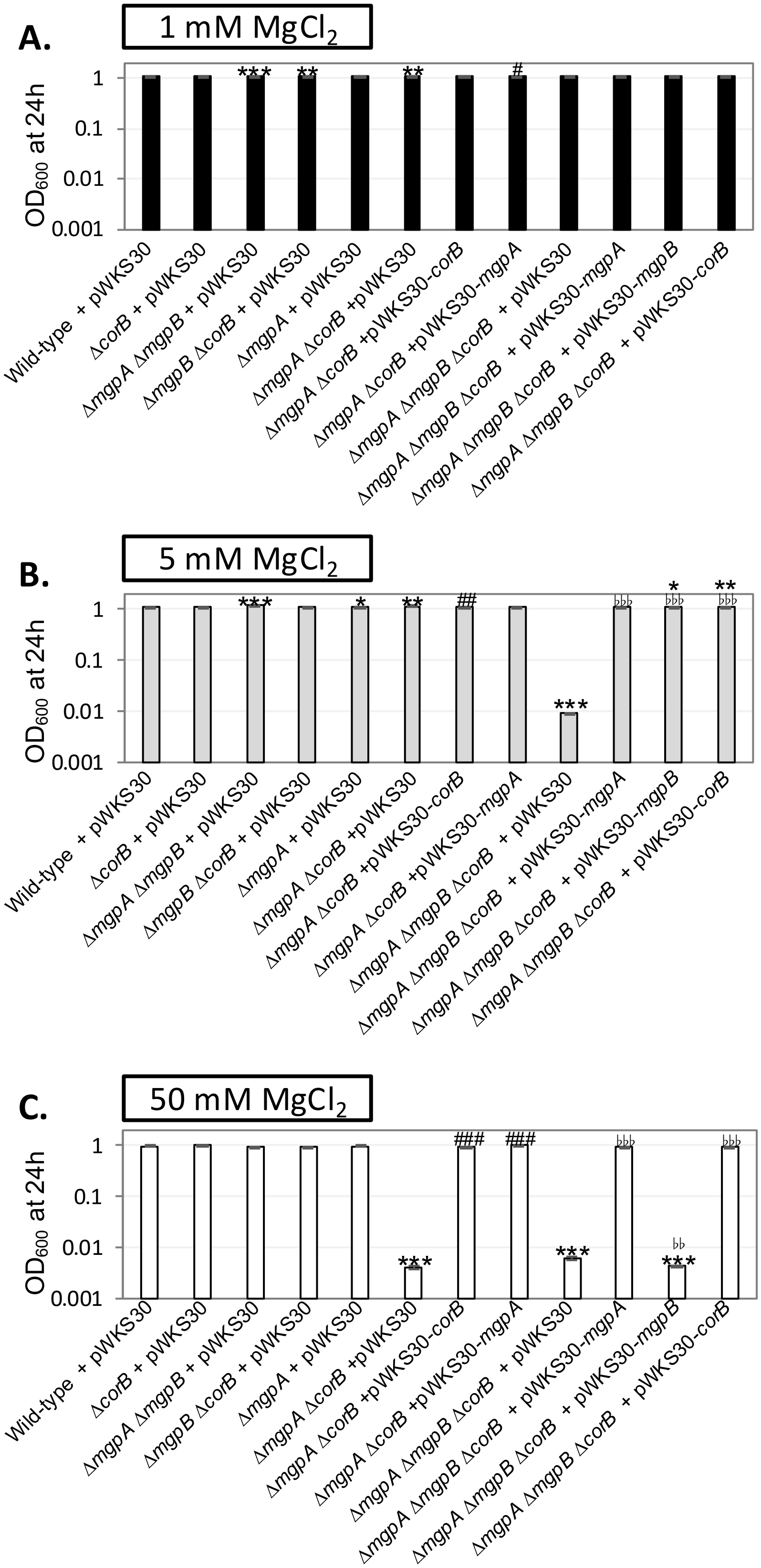


**Fig. S2. Complementation tests in the indicated strains under exogenous high-$Mg^{2+}$ stress at high pH.** The indicated strains were pre-grown to mid-exponential phase in N-minimal medium pH 7.4 with 1 mM $MgCl_2$, washed, and diluted into N-minimal medium pH 8.5 with the indicated amount of $MgCl_2$, followed by incubation at 37 ℃. $OD_{600}$ was measured after 24 hours of incubation. Values are mean ± SD, n = 6. Unpaired t test (p < 0.05*, 0.005**, 0.0005***) versus corresponding Wild-type + pWKS30 , (p < 0.05#, 0.005##, 0.0005###) Δ*mgpA* Δ*corB* +pWKS30 vs Δ*mgpA* Δ*corB* +pWKS30-*corB* or Δ*mgpA* Δ*corB* +pWKS30-*mgpA*, and (p < 0.05 ♭, 0.005 ♭♭, 0.0005 ♭♭♭) Δ*mgpA* Δ*mgpB* Δ*corB* + pWKS30 vs Δ*mgpA* Δ*mgpB* Δ*corB* + pWKS30-*mgpA* or Δ*mgpA* Δ*mgpB* Δ*corB* + pWKS30-*mgpB* or Δ*mgpA* Δ*mgpB* Δ*corB* + pWKS30-*corB*. Strains used: JS2699, JS2768, JS2769, JS2770, JS2771, JS2772, JS2773, JS2774, JS2775, JS2776, JS2777, and JS2778.

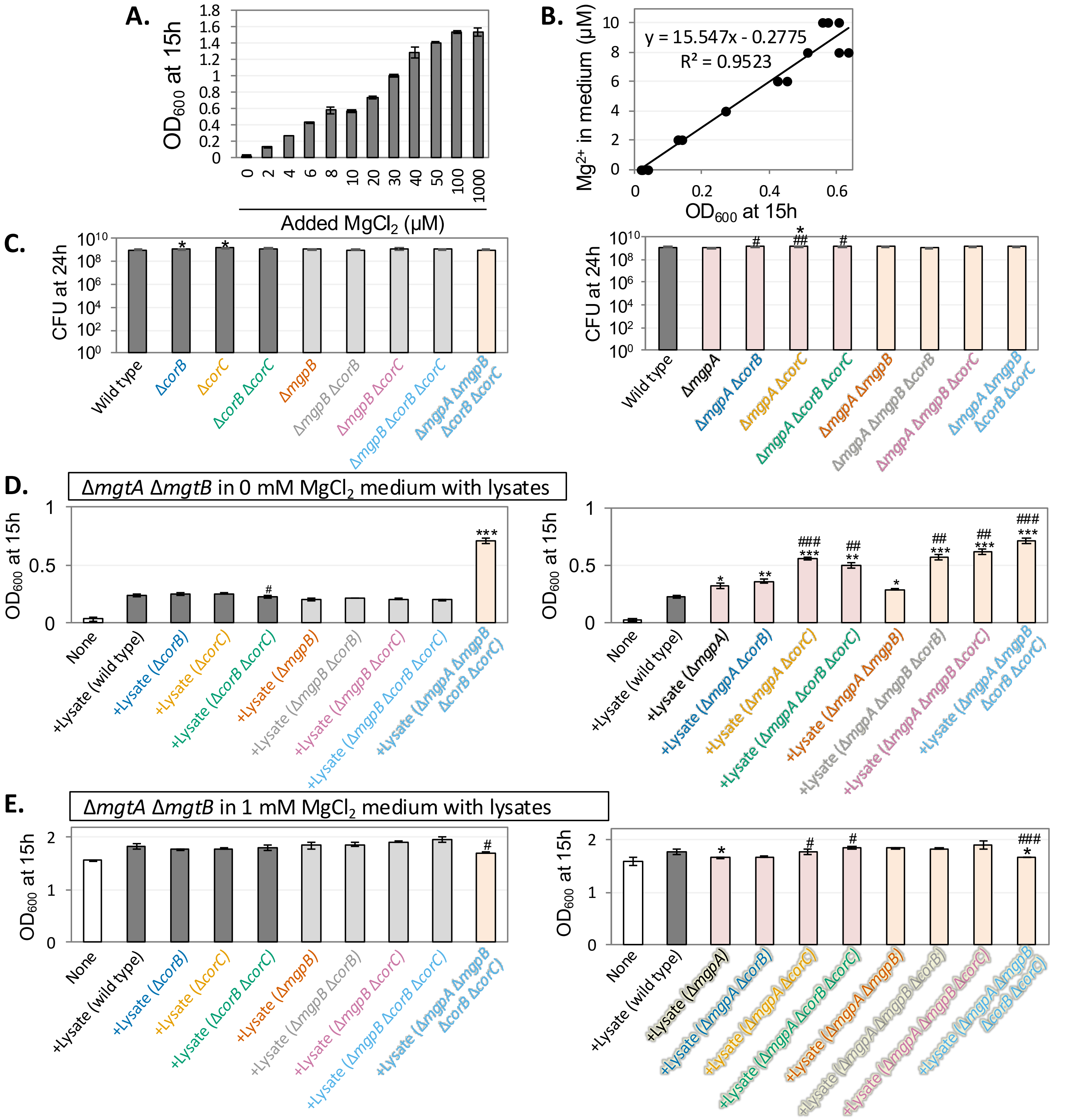


**Fig. S3. The lysate from the Δ*mgpA* Δ*mgpB* Δ*corB* strain grown with $Mg^{2+}$ stimulates the growth of Δ*mgtA* Δ*mgtB* strains in medium without added $Mg^{2+}$ greater than the lysate from the wild type.** The Δ*mgtA* Δ*mgtB* strain was pre-grown to mid-exponential phase in N-minimal medium (pH 7.4) with 10 mM $MgCl_2$ at 37° C, washed, and then diluted into N-minimal medium (pH 7.4) containing the indicated concentrations of $MgCl_2$ and supplemented with 34 μg/mL of the indicated lysate, followed by incubation at 37° C. Lysates were prepared from the indicated strains grown in 10 mM $MgCl_2$, as described in the Materials and Methods. Final $OD_{600}$ values were measured after 15 h of incubation. (A) Growth of the Δ*mgtA* Δ*mgtB* strain in medium containing the indicated concentrations of $MgCl_2$ without lysates. (B) Calibration curve relating final $OD_{600}$ to the $Mg^{2+}$ concentration in the medium, generated from the data in (A). (C) CFU of the indicated strains prior to preparation of the cell lysates. (D) Growth of the Δ*mgtA* Δ*mgtB* strain in N-minimal medium without added $Mg^{2+}$ supplemented with lysates. (E) Growth of the Δ*mgtA* Δ*mgtB* strain in N-minimal medium containing 1 mM $MgCl_2$ supplemented with lysates. Values are mean ± SD, n = 3. Unpaired t-test ($P < 0.05^{*}$, $0.005^{**}$, $0.0005^{***}$) vs. wild type; and ($P < 0.05^{\#}$, $0.005^{\#\#}$, $0.0005^{\#\#\#}$) vs. the corresponding *corB*⁺ *corC*⁺ strains. Strains used: JS2562, 14028, JS2728, JS2740, JS2741, JS2730, JS2729, JS2746, JS2745, JS2748, JS2749, JS2750, JS2751, JS2752, JS2753, JS2754, and JS2755.

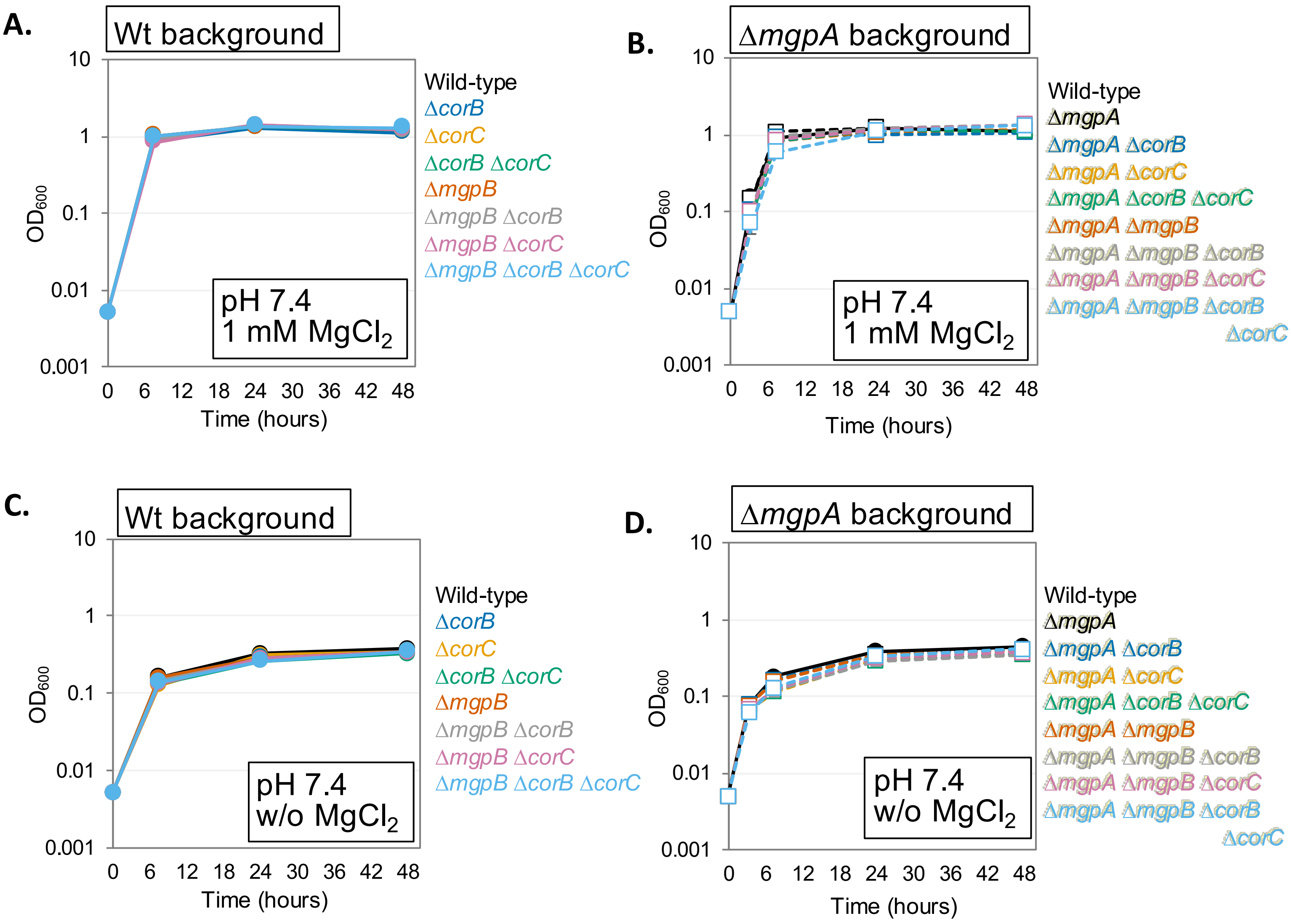


**Fig. S4. $OD_{600}$ of the indicated strains (same experiment as Fig.5) after incubation in $Mg^{2+}$ starvation conditions.** The indicated strains were grown overnight in N-minimal medium pH 7.4 with 1 mM $MgCl_2$, washed, diluted into N-minimal medium pH 7.4 (A and B) with or (C and D) without 1 mM $MgCl_2$ (t = 0 h), and incubated at 37℃. $OD_{600}$ were determined at the indicated time points in (A and C) wild-type and (B and D) Δ*mgpA* background. Values are mean ± SD, n = 6. Unpaired t-test (P < 0.05*, 0.005**, 0.0005***) vs. wild type at the same time point and at the same $Mg^{2+}$ concentration. Complete *p*-values for panels A-D are provided in Tables S10-13, respectively. Strains used: 14028, JS2740, JS2728, JS2741, JS2707, JS2742, JS2743, JS2744, JS2729, JS2745, JS2730, JS2746, JS2747, JS2737, JS2738, and JS2739.

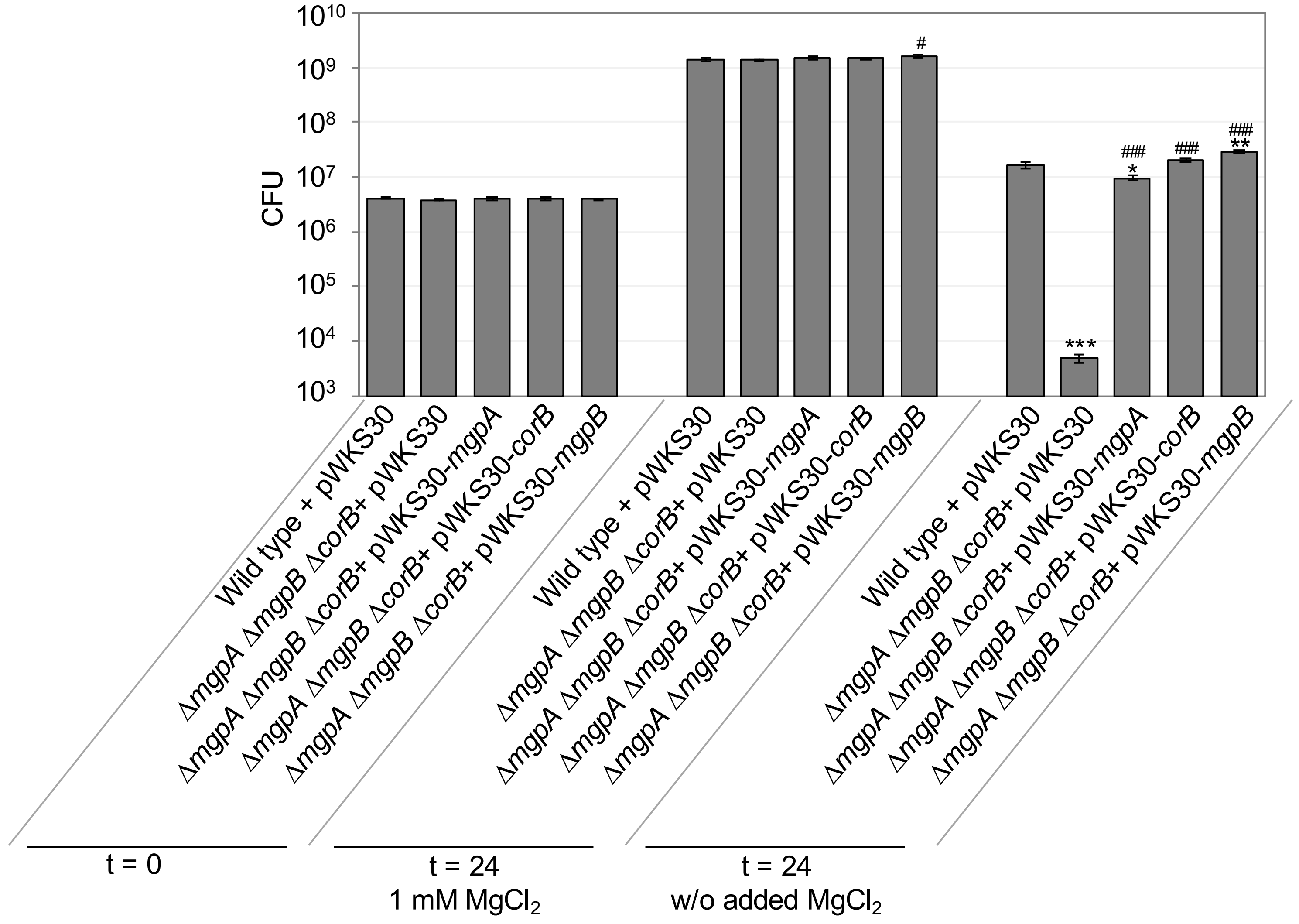


**Fig. S5. Complementation tests in the indicated strains after incubation in $Mg^{2+}$ starvation conditions.** The indicated strains were grown overnight in N-minimal medium pH 7.4 with 1 mM $MgCl_2$, washed, diluted into N-minimal medium pH 7.4 with or without 1 mM $MgCl_2$ (t = 0 h), and incubated at 37℃. CFUs were determined after 24 hours. Values are mean ± SD, n = 6. Unpaired t test ($p < 0.05$*, 0.005**, 0.0005***) versus Wild-type + pWKS30, ($p < 0.05$#, 0.005##, 0.0005###) versus Δ*mgpA* Δ*mgpB* Δ*corB* + pWKS30 at the same time point and at the same $Mg^{2+}$ concentration. Strains used: JS2699, JS2775, JS2776, JS2777, and JS2778.

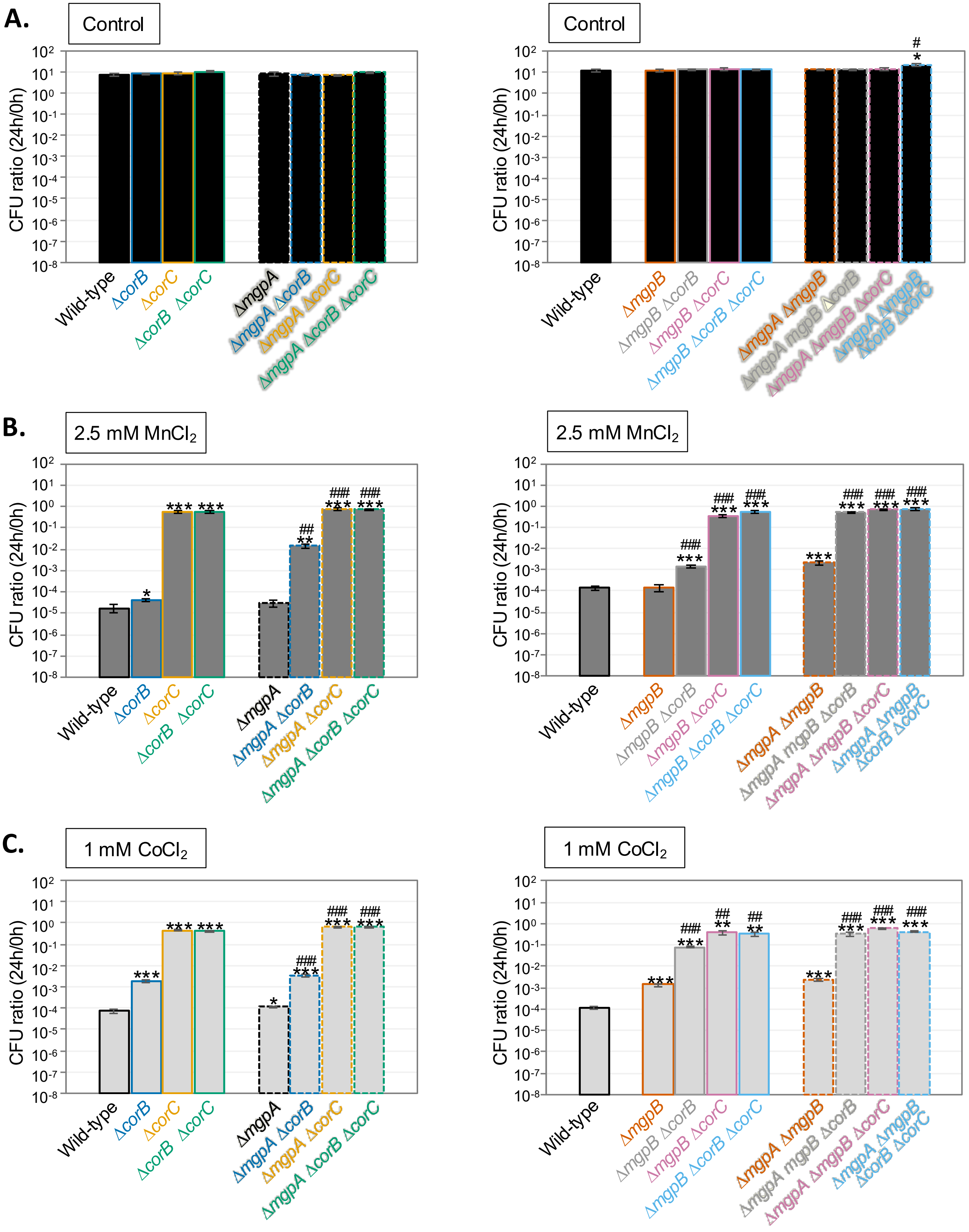


**Fig. S6. Loss of CorB increases tolerance to $Mn^{2+}$ and $Co^{2+}$.** The indicated strains were grown overnight in MOPS-minimal medium (pH 7.4) with 1.32 mM $KH_2PO_4$ and 10 mM $MgCl_2$, washed, diluted into MOPS-minimal medium (pH 7.4) with 50 μM $MgCl_2$ and without $KH_2PO_4$ in the presence or absence of the indicated amounts of $MnCl_2$ or $CoCl_2$ (t = 0 h), and incubated at 37° CFUs were determined at 0 h and 9 h. CFU ratio (9 h/0 h) values are mean ± SD, n = 6. Unpaired t test ($p < 0.05^*$, $0.005^{**}$, $0.0005^{***}$) versus WT, and ($p < 0.05^{\#}$, $0.005^{\#\#}$, $0.0005^{\#\#\#}$) corresponding *corB*$^+$ *corC*. Strains used: 14028, JS2728, JS2740, JS2741, JS2730, JS2729, JS2746, JS2745, JS2748, JS2749, JS2750, JS2751, JS2752, JS2753, JS2754, and JS2755.

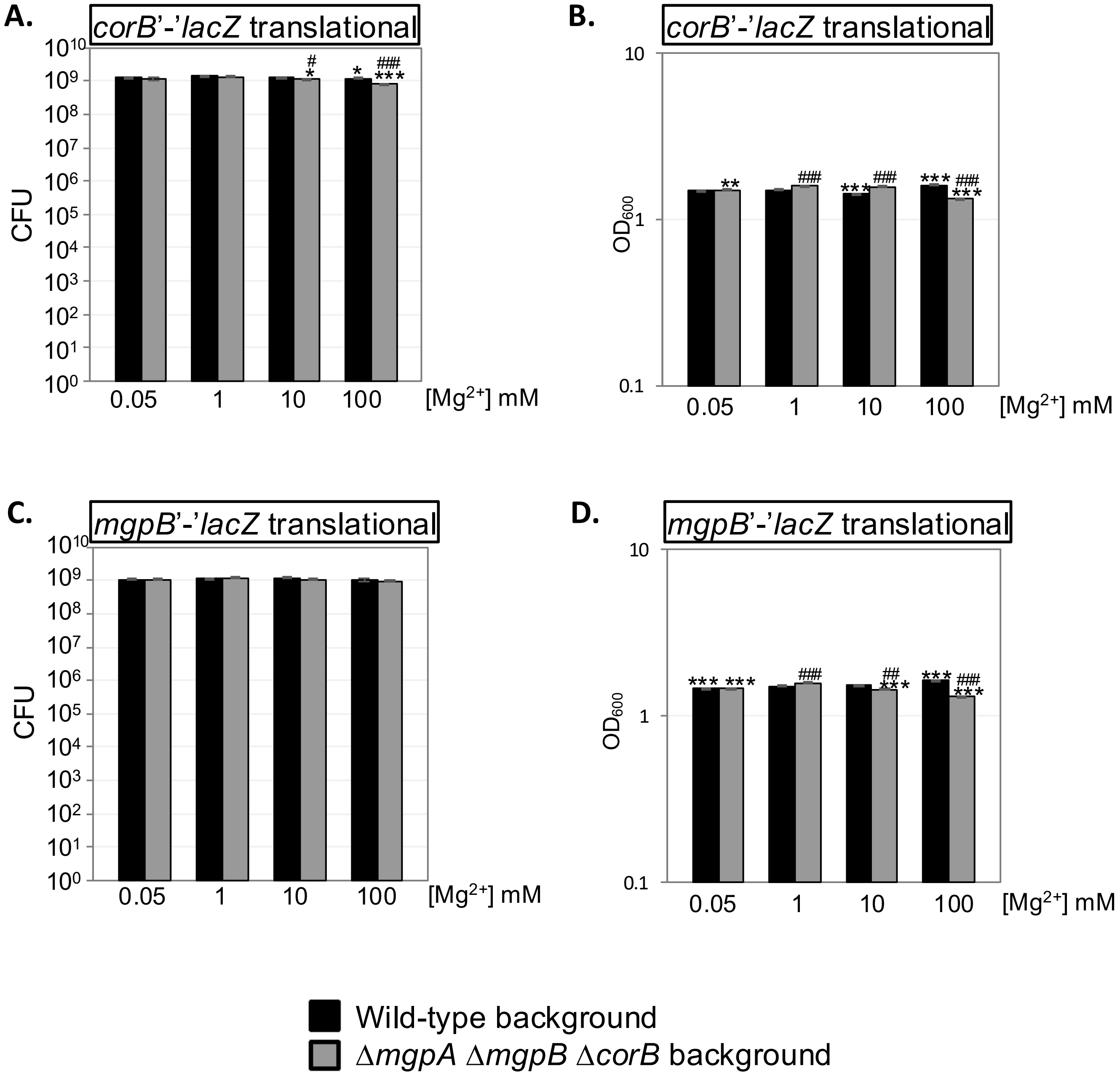


**Fig. S7. Expressions of *corB* and *mgpB* are regulated by $Mg^{2+}$ concentrations.** (A and B) *corB'-'lacZ* and (C and D) *mgpB'-'lacZ* fusions in the wild-type or Δ*mgpA* Δ*mgpB* Δ*corB* backgrounds were pre-grown to mid-exponential phase in N-minimal medium pH 7.4 with 1 mM $MgCl_2$ at 37℃, washed, diluted in N-minimal medium pH 7.4 with or without the indicated amount of $MgCl_2$, and incubated at 37℃ for 24h. (A and C) CFU and (B and D) $OD_{600}$ were determined before β-galactosidase activity was measured. Values are mean ± SD, n = 6. Unpaired t test (p < 0.05*, 0.005**, 0.0005***) versus 1 mM $Mg^{2+}$ in the same background, (p < 0.05#, 0.005##, 0.0005###) vs wild-type background in the same condition. Strains used: JS2764, JS2765, JS2766, and JS2767.

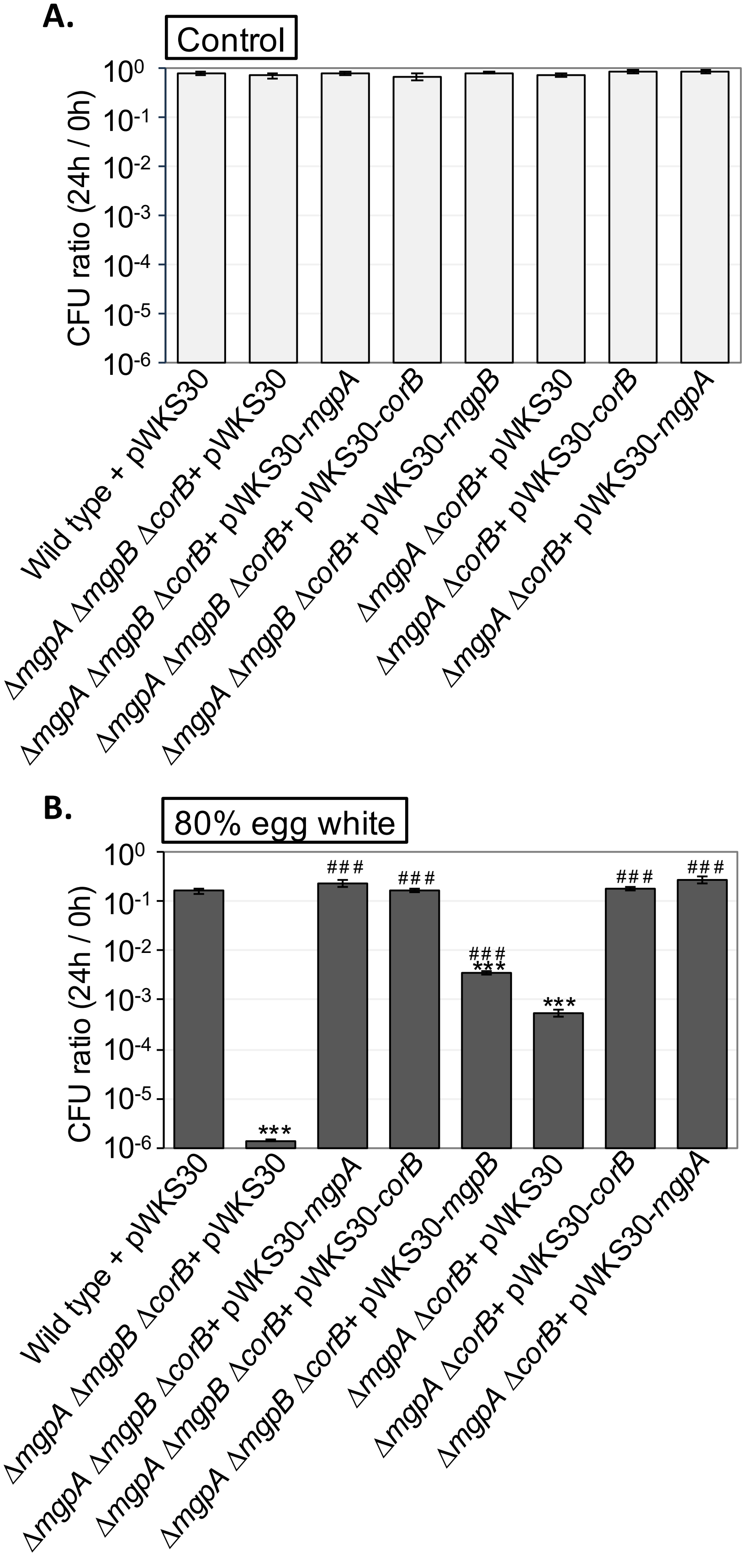


**Fig S8. Complementation tests of egg white tolerance in the indicated strains**. The indicated strains were pre-grown to mid-exponential phase in N-minimal medium (pH 7.4) with 1 mM $MgCl_2$ at 37° C, washed, diluted into (A) PBS buffered with 100 mM HEPES (pH 8.6) or (B) 80% egg white solution (t = 0 h), and incubated at 37° C. CFUs were determined at 0 hour and 24 hours. CFU ratio (24 h/0 h) values are mean ± SD, n = 6. Unpaired t-test (P < 0.05*, 0.005**, 0.0005***) versus WT same treatment, (p < 0.05#, 0.005##, 0.0005###) vs corresponding vector strains. Strains used: JS2699, JS2775, JS2776, JS2777, JS2778, JS2779, JS2772, and JS2774.

**Table S1.** Bacterial strains

| Strain[a] | Genotype | Endpoint[b] | Ref[c] |
|---|---|---|---|
| 14028 | Wild-type | | |
| JS2730 | Δ*mgpA* Δ*corC* | | (1) |
| JS2736 | ΔcorCs6 (Δ*paeA* Δ*corC* Δ*mgpA* Δ*corB* Δ*cvrA* Δ*mgpB*) | | |
| JS2737 | Δ*mgpA* Δ*corB* *mgpB*::Cm | | |
| JS2738 | Δ*mgpA* Δ*corC* *mgpB*::Cm | | |
| JS2739 | Δ*mgpA* Δ*corB* Δ*corC* *mgpB*::Cm | | |
| JS2728 | Δ*corC* | | (1) |
| JS2740 | Δ*corB* | | |
| JS2741 | Δ*corB* Δ*corC* | | |
| JS2707 | *mgpB*::Cm | | (1) |
| JS2742 | Δ*corB* *mgpB*::Cm | | |
| JS2743 | Δ*corC* *mgpB*::Cm | | |
| JS2744 | Δ*corB* Δ*corC* *mgpB*::Cm | | |
| JS2729 | Δ*mgpA* | | (1) |
| JS2745 | Δ*mgpA* Δ*corB* | | |
| JS2746 | Δ*mgpA* Δ*corB* Δ*corC* | | |
| JS2747 | Δ*mgpA* *mgpB*::Cm | | |
| JS2562 | Δ*mgtA*::Kan Δ*mgtB* | | (2) |
| JS2748 | Δ*mgpA* Δ*mgpB* | | |
| JS2749 | Δ*mgpA* Δ*mgpB* Δ*corB* | | |
| JS2750 | Δ*mgpA* Δ*mgpB* Δ*corC* | | |
| JS2751 | Δ*mgpA* Δ*mgpB* Δ*corB* Δ*corC* | | |
| JS2752 | Δ*mgpB* | | |
| JS2753 | Δ*mgpB* Δ*corB* | | |
| JS2754 | Δ*mgpB* Δ*corC* | | |
| JS2755 | Δ*mgpB* Δ*corB* Δ*corC* | | |
| JS2560 | Δ*speED* Δ*speG* Δ*speF* Δ*cadA* Δ*speC* Δ*ldcC* Δ*speB* Δ*speA*::Tc (Δsynth) | | (2) |
| JS2756 | Δsynth Δ*mgpA* Δ*corB* *mgpB*::Cm | | |
| JS2713 | Δ*mgtA* Δ*mgtB* | | (1) |

| | | | |
|---|---|---|---|
| JS2757 | Δ*mgtA* Δ*mgtB* Δ*mgpA* Δ*corB* *mgpB*::Cm | | |
| JS2720 | Δ*corA* | | (1) |
| JS2758 | Δ*corA* Δ*mgpA* Δ*mgpB* Δ*corB* | | |
| JS2695 | Δ*paeA* | | (1) |
| JS2759 | Δ*paeA* Δ*mgpA* Δ*mgpB* Δ*corB* | | |
| JS2760 | Δ*corD* | 103000-103369 | |
| JS2761 | Δ*mgpA* Δ*mgpB* Δ*corB* Δ*corD* | | |
| JS2762 | Δ*corA* Δ*corD* | | |
| JS2763 | Δ*corA* Δ*mgpA* Δ*mgpB* Δ*corB* Δ*corD* | | |
| JS2764 | att::*corB*-translation-lacZ | 2871646-2873465 | |
| JS2765 | Δ*mgpA* Δ*mgpB* Δ*corB* att::*corB*-translation-lacZ | | |
| JS2766 | att::*mgpB*1 translation-lacZ | 2253911-2254880 | |
| JS2767 | Δ*mgpA* Δ*mgpB* Δ*corB* att::*mgpB*1 translation-lacZ | | |
| JS2693 | Δ*mgpA*::Cm | | (1) |
| JS2699 | Wild-type + pWKS30 | | (1) |
| JS2768 | Δ*corB* + pWKS30 | | |
| JS2769 | Δ*mgpA* Δ*mgpB* + pWKS30 | | |
| JS2770 | Δ*mgpB* Δ*corB* + pWKS30 | | |
| JS2771 | Δ*mgpA* + pWKS30 | | |
| JS2772 | Δ*mgpA* Δ*corB* +pWKS30 | | |
| JS2773 | Δ*mgpA* Δ*corB* + pWKS30-placZ-*corB* | | |
| JS2774 | Δ*mgpA* Δ*corB* + pWKS30-*mgpA* | | |
| JS2775 | Δ*mgpA* Δ*mgpB* Δ*corB* + pWKS30 | | |
| JS2776 | Δ*mgpA* Δ*mgpB* Δ*corB* + pWKS30-*mgpA* | | |
| JS2777 | Δ*mgpA* Δ*mgpB* Δ*corB* + pWKS30-placZ-*corB* | | |
| JS2778 | Δ*mgpA* Δ*mgpB* Δ*corB* + pWKS30-*mgpB* | | |
| JS2779 | Δ*paeA* Δ*corB* | | |
| JS2724 | Δ*paeA* Δ*corC* | | (1) |
| JS2780 | Δ*paeA* Δ*corB* Δ*corC* | | |
| JS2781 | Δ*paeA* Δ*mgpA* Δ*mgpB* | | |
| JS2782 | Δ*paeA* Δ*mgpA* Δ*mgpB* Δ*corC* | | |
| JS2783 | ΔpaeA Δ*mgpA* Δ*mgpB* Δ*corB* Δ*corC* | | |

a All *Salmonella* strains are isogenic derivatives of *S. enterica* serovar Typhimurium strain 14028.

b Numbers indicate the base pairs that are deleted (inclusive) as defined in the *S. enterica* serovar Typhimurium 14028 genome sequence (National Center for Biotechnology Information; NC_016856.1)
c This study, unless otherwise indicated

**Table 2.** Plasmids

| Name | Characteristic | Cloned endpoint[a] | Ref[b] |
|---|---|---|---|
| pKD46 | *bla* PBAD *gam bet exo* pSC101 *oriTS* | | (3) |
| pCP20 | *bla cat* cI857 λPR *flp* pSC101 *oriTS* | | (3) |
| pKD3 | *bla* FRT *cat* FRT PS1 PS2 *oriR6K* | | (3) |
| pKD4 | *bla* FRT *aph* FRT PS1 PS2 *oriR6K* | | (3) |
| pWKS30 | *bla* pSC101 *ori* | | (4) |
| pWKS30-placZ-*corB* | | 2873457-2874746 | |
| pWKS30-*mgpB* | | 2254204-2256543 | |
| pWKS30-*mgpA* | | | (1) |
| pDX1 | *lacZ* tL3 λ*attP oriR6K aacIV tmgB* | | (5) |
| pDX1-*corB*-translation-lacZ | | 2871646-2873465 | |
| pDX1-*mgpB*1 translation-lacZ | | 2253911-2254880 | |

a Numbers indicate the base pairs that are cloned (inclusive) as defined in the *S. enterica* serovar Typhimurium 14028 genome sequence (National Center for Biotechnology Information; NC_016856.1)
b This study, unless otherwise indicated

**Table S3.** Primers used

| Name | Sequence (5' to 3') | Usage |
|---|---|---|
| YIV_pWKS30-10 | AGCTGTTTCCTGTGTGAAATTGTTATCC | pWKS30-placZ-*corB* |
| YIV_pWKS30-12 | ATTGCGTTGCGCTCACTGCC | pWKS30-mgpB |
| YIV_tR2N | AACAGGCCTGCTGGTAATCG | pWKS30-placZ-c*orB,* pWKS30-*mgpB* |
| YIS_STM-corBC | ACCAGCAGGCCTGTTGTGTTACTCCGCCACACTCTC | pWKS30-placZ-*corB* |
| YIS_RplacZC-STM-corBN | ACACAGGAAACAGCTCTGGAACACATCTCCACCAC | pWKS30-placZ-*corB* |
| YIS_225-7 | TGAGCGCAACGCAATTGATTATTTATCACTTTGGCAGAGTCATTATC | pWKS30-*mgpB* |
| YIS_225-9 | ACCAGCAGGCCTGTTTTTTTGGCCAAAGGGCAGAG | pWKS30-*mgpB* |
| YIV_pDX1-18 | GCAAGCTTGGCACTGGCCAC | pDX1-*corB*-translation-lacZ, pDX1-*mgpB*1 translation-lacZ |
| YIV_pDX1-19 | ACCATGATTACGGATTCACTG | pDX1-*corB*-translation-lacZ, pDX1-*mgpB*1 translation-lacZ |
| YIS_287-20 | CAGTGCCAAGCTTGCTTTTCTCGCCGACACCGAG | pDX1-*corB*-translation-lacZ |
| YIS_287-21 | ATCCGTAATCATGGTGTGTTCCAGGGGAAACTCCTTTAAAAG | pDX1-*corB*-translation-lacZ |
| YIS_225-10 | CAGTGCCAAGCTTGCCAGACGGTGACATTCAGCAC | pDX1-*mgpB*1 translation-*lacZ* |
| YIS_225-18 | ATCCGTAATCATGGTCATAGTAAGAGCAATACCTTCTGTAATATGACAATTG | pDX1-*mgpB1 translation-lacZ* |

**Table S4.** P-values for Fig. 3B. Unpaired t test versus corresponding WT at the same condition. Red cells indicate $p < 0.0005$, and yellow cells indicate $p < 0.005$.

| | 1 mM | 25 mM | 100 mM | 250 mM |
|---|---|---|---|---|
| Δ*corC* | 4.81E-03 | 3.22E-03 | 3.22E-03 | 8.38E-07 |
| Δ*corB* | 1.94E-03 | 8.09E-02 | 8.09E-02 | 1.23E-01 |
| Δ*mgpB* | 2.70E-04 | 2.11E-01 | 2.11E-01 | 2.76E-01 |
| Δ*corB* Δ*corC* | 3.34E-04 | 5.00E-01 | 5.00E-01 | 3.28E-05 |
| Δ*mgpB* Δ*corB* | 1.04E-03 | 5.76E-02 | 5.76E-02 | 4.70E-01 |
| Δ*mgpB* Δ*corC* | 1.75E-03 | 5.00E-01 | 5.00E-01 | 2.27E-05 |
| Δ*mgpB* Δ*corB* Δ*corC* | 5.15E-04 | 7.89E-02 | 7.89E-02 | 3.23E-05 |
| Δ*mgpA* | 6.86E-04 | 2.47E-01 | 2.47E-01 | 1.09E-07 |
| Δ*mgpA* Δ*mgpB* | 1.32E-04 | 7.36E-04 | 7.36E-04 | 8.06E-06 |

**Table S5.** P-values for Fig. 3C. Unpaired t test versus corresponding WT at the same condition. Red cells indicate $p < 0.0005$, yellow cells indicate $p < 0.005$, and green cells indicate $p < 0.005$.

| | 1 mM | 2.5 mM | 5 mM | 25 mM |
|---|---|---|---|---|
| Δ*mgpA* | 5.24E-02 | 1.05E-01 | 7.98E-02 | 4.38E-01 |
| Δ*mgpA* Δ*corC* | 5.64E-02 | 2.42E-01 | 1.05E-10 | 2.76E-16 |
| Δ*mgpA* Δ*corB* | 3.24E-01 | 2.65E-02 | 5.00E-01 | 1.86E-16 |
| Δ*mgpA* Δ*mgpB* | 1.41E-02 | 4.14E-01 | 1.14E-01 | 3.15E-01 |
| Δ*mgpA* Δ*corB* Δ*corC* | 4.23E-01 | 4.28E-01 | 3.09E-13 | 1.84E-16 |
| Δ*mgpA* Δ*mgpB* Δ*corB* | 1.03E-01 | 4.62E-10 | 1.70E-18 | 1.85E-16 |
| Δ*mgpA* Δ*mgpB* Δ*corC* | 1.61E-01 | 1.42E-16 | 1.69E-18 | 1.86E-16 |
| Δ*mgpA* Δ*mgpB* Δ*corB* Δ*corC* | 2.16E-03 | 7.07E-13 | 3.17E-18 | 1.86E-16 |

**Table S6.** P-values for Fig. 5A. Unpaired t-test vs. wild type at the same time point and at the same $Mg^{2+}$ concentration. Yellow cells indicate p < 0.005 and green cells indicate p < 0.005.

| | t=0 | t=7.5 | t=24 | t=48 |
|---|---|---|---|---|
| *ΔcorC* | 9.69E-01 | 6.48E-01 | 1.00E+00 | 9.48E-02 |
| *ΔcorB* | 9.49E-01 | 7.91E-01 | 3.03E-01 | 1.07E-01 |
| *ΔmgpB* | 3.90E-01 | 3.21E-02 | 6.21E-01 | 1.86E-01 |
| *ΔcorC ΔcorB* | 6.47E-02 | 7.43E-01 | 8.98E-02 | 8.06E-03 |
| *ΔcorB ΔmgpB* | 8.08E-01 | 1.59E-01 | 8.30E-01 | 2.12E-02 |
| *ΔcorC ΔmgpB* | 8.71E-01 | 3.04E-01 | 5.23E-01 | 9.46E-04 |
| *ΔcorC ΔcorB ΔmgpB* | 5.41E-01 | 5.52E-01 | 5.95E-01 | 6.00E-02 |

**Table S7.** P-values for Fig. 5B. Unpaired t-test (P < 0.05*, 0.005**, 0.0005***) vs. wild type at the same time point and at the same $Mg^{2+}$ concentration. Red cells indicate p < 0.0005, yellow cells indicate p < 0.005, and green cells indicate p < 0.005.

| | t=0 | t=3.5 | t=7.5 | t=24 | t=48 |
|---|---|---|---|---|---|
| *ΔmgpA* | 8.24E-01 | 2.35E-01 | 9.54E-02 | 8.62E-01 | 4.96E-02 |
| *ΔmgpA ΔcorC* | 1.55E-01 | 5.02E-01 | 5.22E-01 | 4.93E-01 | 2.27E-03 |
| *ΔmgpA ΔcorB* | 3.57E-01 | 9.41E-01 | 1.26E-01 | 6.50E-01 | 3.79E-02 |
| *ΔmgpA ΔmgpB* | 4.89E-03 | 3.24E-01 | 1.86E-01 | 6.07E-01 | 8.38E-05 |
| *ΔmgpA ΔcorC ΔcorB* | 6.42E-03 | 2.39E-01 | 8.73E-01 | 8.50E-01 | 7.33E-04 |
| *ΔmgpA ΔcorB ΔmgpB* | 2.94E-02 | 7.43E-01 | 4.99E-02 | 2.09E-01 | 4.58E-04 |
| *ΔmgpA ΔcorC ΔmgpB* | 2.28E-03 | 5.22E-01 | 1.04E-01 | 7.24E-01 | 1.18E-02 |
| *ΔmgpA ΔcorC ΔcorB ΔmgpB* | 2.64E-01 | 1.24E-02 | 6.69E-01 | 8.04E-01 | 1.08E-01 |

**Table S8.** P-values for Fig. 5C. Unpaired t-test (P < 0.05*, 0.005**, 0.0005***) vs. wild type at the same time point and at the same $Mg^{2+}$ concentration. Red cells indicate p < 0.0005, yellow cells indicate p < 0.005, and green cells indicate p < 0.005.

| | t=0 | t=7.5 | t=24 | t=48 |
|---|---|---|---|---|
| Δ*corC* | 9.69E-01 | 3.40E-02 | 8.99E-06 | 1.27E-04 |
| Δ*corB* | 9.49E-01 | 9.06E-01 | 6.07E-01 | 9.27E-01 |
| Δ*mgpB* | 3.90E-01 | 4.63E-01 | 3.82E-01 | 9.04E-01 |
| Δ*corC* Δ*corB* | 6.47E-02 | 5.37E-03 | 4.45E-06 | 1.22E-04 |
| Δ*corB* Δ*mgpB* | 8.08E-01 | 1.35E-02 | 6.15E-01 | 7.52E-01 |
| Δ*corC* Δ*mgpB* | 8.71E-01 | 1.36E-05 | 7.48E-06 | 1.16E-04 |
| Δ*corC* Δ*corB* Δ*mgpB* | 5.41E-01 | 1.46E-03 | 6.47E-06 | 1.73E-03 |

**Table S9.** P-values for Fig. 5D. Unpaired t-test (P < 0.05*, 0.005**, 0.0005***) vs. wild type at the same time point and at the same $Mg^{2+}$ concentration. Red cells indicate p < 0.0005, yellow cells indicate p < 0.005, and green cells indicate p < 0.005.

| | t=0 | t=3.5 | t=7.5 | t=24 | t=48 |
|---|---|---|---|---|---|
| Δ*mgpA* | 8.24E-01 | 4.59E-01 | 4.06E-01 | 4.20E-01 | 8.83E-01 |
| Δ*mgpA* Δ*corC* | 1.55E-01 | 1.05E-01 | 1.17E-06 | 3.67E-09 | 2.20E-06 |
| Δ*mgpA* Δ*corB* | 3.57E-01 | 2.25E-01 | 5.60E-01 | 9.13E-01 | 7.09E-01 |
| Δ*mgpA* Δ*mgpB* | 4.89E-03 | 1.10E-01 | 3.88E-01 | 8.01E-01 | 2.28E-01 |
| Δ*mgpA* Δ*corC* Δ*corB* | 6.42E-03 | 1.33E-02 | 1.13E-06 | 3.67E-09 | 2.20E-06 |
| Δ*mgpA* Δ*corB* Δ*mgpB* | 2.94E-02 | 5.10E-02 | 1.18E-06 | 3.67E-09 | 2.20E-06 |
| Δ*mgpA* Δ*corC* Δ*mgpB* | 2.28E-03 | 1.46E-02 | 1.20E-06 | 3.67E-09 | 2.03E-04 |
| Δ*mgpA* Δ*corC* Δ*corB* Δ*mgpB* | 2.64E-01 | 6.65E-03 | 1.12E-06 | 3.67E-09 | 2.20E-06 |

**Table S10.** P-values for Fig. S4A. Values were calculated against the wild type under the same conditions. Red cells indicate $p < 0.0005$, yellow cells indicate $p < 0.005$, and green cells indicate $p < 0.005$.

| | t=7.5 | t=24 | t=48 |
|---|---|---|---|
| Δ*corC* | 1.56E-01 | 9.20E-01 | 4.75E-01 |
| Δ*corB* | 1.70E-01 | 3.72E-03 | 9.25E-01 |
| Δ*mgpB* | 6.55E-01 | 4.83E-01 | 6.65E-03 |
| Δ*corC* Δ*corB* | 8.04E-01 | 2.98E-01 | 8.88E-02 |
| Δ*corB* Δ*mgpB* | 3.53E-01 | 1.19E-01 | 1.40E-03 |
| Δ*corC* Δ*mgpB* | 5.30E-02 | 1.78E-03 | 1.22E-04 |
| Δ*corC* Δ*corB* Δ*mgpB* | 9.20E-01 | 3.04E-01 | 3.14E-03 |

**Table S11.** P-values for Fig. S4B. Values were calculated against the wild type under the same conditions. Red cells indicate $p < 0.0005$, yellow cells indicate $p < 0.005$, and green cells indicate $p < 0.005$.

| | t=3.5 | t=7.5 | t=24 | t=48 |
|---|---|---|---|---|
| Δ*mgpA* | 9.13E-01 | 1.44E-01 | 9.61E-01 | 1.00E+00 |
| Δ*mgpA* Δ*corC* | 6.32E-02 | 9.20E-01 | 2.26E-03 | 7.72E-01 |
| Δ*mgpA* Δ*corB* | 9.64E-02 | 8.83E-01 | 1.58E-04 | 7.06E-02 |
| Δ*mgpA* Δ*mgpB* | 6.81E-02 | 4.22E-01 | 1.29E-02 | 2.66E-01 |
| Δ*mgpA* Δ*corC* Δ*corB* | 5.88E-02 | 3.45E-01 | 8.53E-02 | 5.69E-01 |
| Δ*mgpA* Δ*corB* Δ*mgpB* | 1.84E-01 | 9.06E-01 | 5.14E-01 | 1.70E-04 |
| Δ*mgpA* Δ*corC* Δ*mgpB* | 1.12E-01 | 5.82E-01 | 1.40E-01 | 6.64E-05 |
| Δ*mgpA* Δ*corC* Δ*corB* Δ*mgpB* | 2.39E-02 | 2.18E-02 | 2.84E-02 | 4.30E-06 |

**Table S12** P-values for Fig. S4C. Values were calculated against the wild type under the same conditions. Yellow cells indicate p < 0.005 and green cells indicate p < 0.005.

| | t=7.5 | t=24 | t=48 |
|---|---|---|---|
| Δ*corC* | 6.71E-03 | 5.49E-01 | 4.22E-01 |
| Δ*corB* | 3.70E-01 | 1.52E-01 | 2.53E-01 |
| Δ*mgpB* | 5.93E-01 | 1.40E-01 | 2.76E-01 |
| Δ*corC* Δ*corB* | 9.81E-03 | 4.06E-02 | 1.66E-01 |
| Δ*corB* Δ*mgpB* | 1.06E-03 | 9.11E-02 | 3.04E-01 |
| Δ*corC* Δ*yegH* | 6.32E-04 | 1.17E-01 | 2.22E-01 |
| Δ*corC* Δ*corB* Δ*mgpB* | 8.51E-02 | 4.11E-02 | 3.69E-01 |

**Table S13.** P-values for Fig. S4D. Values were calculated against the wild type under the same conditions. Yellow cells indicate p < 0.005 and green cells indicate p < 0.005.

| | t=3.5 | t=7.5 | t=24 | t=48 |
|---|---|---|---|---|
| Δ*mgpA* | 9.42E-01 | 2.64E-01 | 1.19E-01 | 2.51E-01 |
| Δ*mgpA* Δ*corC* | 2.65E-01 | 4.78E-03 | 2.43E-03 | 6.93E-03 |
| Δ*mgpA* Δ*corB* | 1.00E+00 | 3.37E-01 | 5.60E-02 | 1.76E-01 |
| Δ*mgpA* Δ*mgpB* | 8.82E-01 | 1.80E-01 | 3.98E-02 | 1.02E-01 |
| Δ*mgpA* Δ*corC* Δ*corB* | 2.50E-01 | 1.73E-03 | 1.33E-03 | 1.04E-02 |
| Δ*mgpA* Δ*corB* Δ*mgpB* | 5.55E-01 | 2.13E-02 | 3.92E-03 | 7.34E-03 |
| Δ*mgpA* Δ*corC* Δ*mgpB* | 2.98E-01 | 1.44E-02 | 1.52E-02 | 4.14E-02 |
| Δ*mgpA* Δ*corC* Δ*corB* Δ*mgpB* | 1.43E-01 | 3.10E-02 | 6.21E-02 | 3.28E-01 |

**Table S14.** P-values for Fig. S6A-F. Values were calculated by comparing the Δ*mgpA* Δ*mgpB* Δ*corB* strain with the *mgpA*⁺ *mgpB*⁺ *corB*⁺ strain in the same background under the same conditions. Red cells indicate p < 0.0005, yellow cells indicate p < 0.005, and green cells indicate p < 0.005.

| | $Mg^{2+}$ | t=0 | t=3.5 | t=7.5 | t=24 | t=48 |
|---|---|---|---|---|---|---|
| Δsynth | + | 4.11E-01 | 2.20E-03 | 4.88E-05 | 9.75E-01 | 8.45E-01 |
| | - | 4.11E-01 | 8.81E-03 | 1.59E-08 | 9.83E-07 | 1.18E-05 |
| Δ*mgtAB* | + | 3.89E-01 | 2.75E-03 | 4.27E-04 | 9.05E-01 | 6.87E-01 |
| | - | 3.89E-01 | 8.72E-02 | 3.16E-02 | 6.18E-01 | 5.59E-02 |
| Δ*corA* | + | 1.17E-02 | - | 2.45E-01 | 3.39E-02 | 4.90E-02 |
| | - | 1.17E-02 | 3.17E-07 | 5.61E-07 | 2.44E-04 | 7.97E-04 |
| Δ*paeA* | + | 3.29E-01 | - | 8.42E-01 | 4.04E-01 | 8.56E-03 |
| | - | 3.29E-01 | 2.36E-05 | 2.35E-06 | 8.01E-02 | 2.68E-05 |
| Δ*corA*Δ*corD* | + | 9.51E-03 | - | 7.39E-02 | 9.46E-02 | 6.41E-01 |
| | - | 9.51E-03 | 2.34E-03 | 5.18E-09 | 4.49E-09 | 2.07E-08 |
| Δ*corD* | + | 6.18E-01 | - | 4.55E-01 | 9.61E-01 | 9.55E-01 |
| | - | 6.18E-01 | 6.16E-02 | 2.24E-07 | 1.53E-07 | 2.29E-07 |
